# Genomic signals found using RNA sequencing support conservation of walleye (*Sander vitreus*) in a large freshwater ecosystem

**DOI:** 10.1101/2020.02.07.937961

**Authors:** Matt J. Thorstensen, Jennifer D. Jeffrey, Jason R. Treberg, Douglas A. Watkinson, Eva C. Enders, Ken M. Jeffries

## Abstract

RNA sequencing is an effective approach for studying an aquatic species with little prior molecular information available, yielding both physiological and genomic data, but its genetic applications are not well-characterized. We investigate this possible role for RNA sequencing for population genomics in Lake Winnipeg, Manitoba, Canada, walleye (*Sander vitreus*). Lake Winnipeg walleye represent the largest component of the second-largest freshwater fishery in Canada. In the present study, large female walleye were sampled via nonlethal gill biopsy over two years at three spawning sites representing a latitudinal gradient in the lake. Genetic variation from sequenced messenger RNA was analyzed for neutral and adaptive markers to investigate population structure and possible adaptive variation. We find low population divergence (*F*_ST_ = 0.0095), possible northward gene flow, and outlier loci that vary latitudinally in transcripts associated with cell membrane proteins and cytoskeletal function. These results indicate that Lake Winnipeg walleye may be effectively managed as a single demographically connected metapopulation with contributing subpopulations, and suggest genomic differences possibly underlying observed phenotypic differences. Because RNA sequencing data can yield physiological in addition to genetic information discussed here, we argue that it is useful for addressing diverse molecular questions in the conservation of freshwater species.

## Introduction

Population abundances in aquatic systems are in decline globally, with a 36% decline in the marine Living Planet Index (LPI, http://livingplanetindex.org) between 1970 and 2012, and an 81% decline in the freshwater LPI during the same period (WWF 2016). These estimates are especially alarming for freshwater ecosystems, which cover 2.3% of the earth’s global land surface area but are disproportionately high in species richness—for instance, one-third of all described vertebrate species live in freshwater (Reid et al. 2019; WWF 2018). It is therefore a significant concern that freshwater species are declining in abundance more rapidly than terrestrial or marine species (Reid et al. 2019). This decline underscores an urgent need for research supporting conservation efforts for these diverse freshwater species.

To take effective action, conservation practitioners require research on the environmental stressors a population faces, as well as population structure and evolutionary patterns to determine a species’ adaptive potential (Connon et al. 2018; Russello et al. 2011; Waples & Gaggiotti 2006). Transcriptomics has been discussed in the context of differential gene expression, for identifying important physiological thresholds in species of conservation concern that can support risk assessments and setting management thresholds, thus, ultimately benefiting species conservation (Connon et al. 2018). An advantage of using RNA sequencing for conservation research is that it provides information about both genetics and molecular physiology by returning transcript abundances and single nucleotide polymorphisms (SNPs) allowing researchers to gather a diverse array of information within one data set. These advantages make transcriptomics approaches useful for studying species of conservation concern, especially for species that do not have extensive molecular databases like those available for model species (e.g., zebrafish, *Danio rerio*).

Applications for transcriptomics to address population genomics questions in non-model species is relatively poorly characterized. A large topic of interest in conservation genomics is population structure, or genomic divergence between different groups of individuals, which can support decisions on whether those groups should be managed as a single or several units (Funk et al. 2012). Complementary to population structure analyses, outlier SNP detection may reveal adaptive variation useful for conservation (Funk et al. 2012; Russello et al. 2011). While many genomic methods rely on genes in linkage with outlier SNPs of interest to interpret the functional significance of data, the functional significance of SNPs in mRNA is more readily interpretable because those SNPs may represent *cis*-regulatory mechanisms within an annotated transcript (Verta & Jones 2019). Within a transcript, the effects of SNPs in open reading frames can be predicted, indicating how protein function may be modified by genetic variation (Cingolani et al 2012). Therefore, RNA sequencing may be an effective method for characterizing physiological patterns, population structure, and adaptive variation in species and systems with little prior information available.

Walleye (*Sander vitreus*) in Lake Winnipeg, Manitoba, are the largest component of the second largest freshwater fishery in Canada. Lake Winnipeg is characterized by a north and a south basin connected by a narrow channel (Johnston et al. 2012; Figure 1). While previous microsatellite research showed slight population differentiation between groups in each basin (Backhouse-James & Docker 2011), morphological, life history, dietary, and environmental differences among Lake Winnipeg walleye suggest diverging genetic histories (Environment Canada 2011; Johnson et al. 2012; Moles et al. 2010; Sheppard et al. 2015, Sheppard et al. 2018; Watkinson & Gillis 2005). Within Lake Winnipeg, walleye have shown declining biomass and body condition, decreased catches, and commercial harvests above maximum sustainable yields for several years (Manitoba Government 2018; Manitoba Sustainable Development 2018). Observations of dwarf walleye suggest signs of selection against large, economically desirable fish (Johnston et al. 2012; Moles et al. 2010). These trends highlight the need to gain information on population structure and biological differences in Lake Winnipeg walleye to support future conservation efforts.

**Figure 1.**
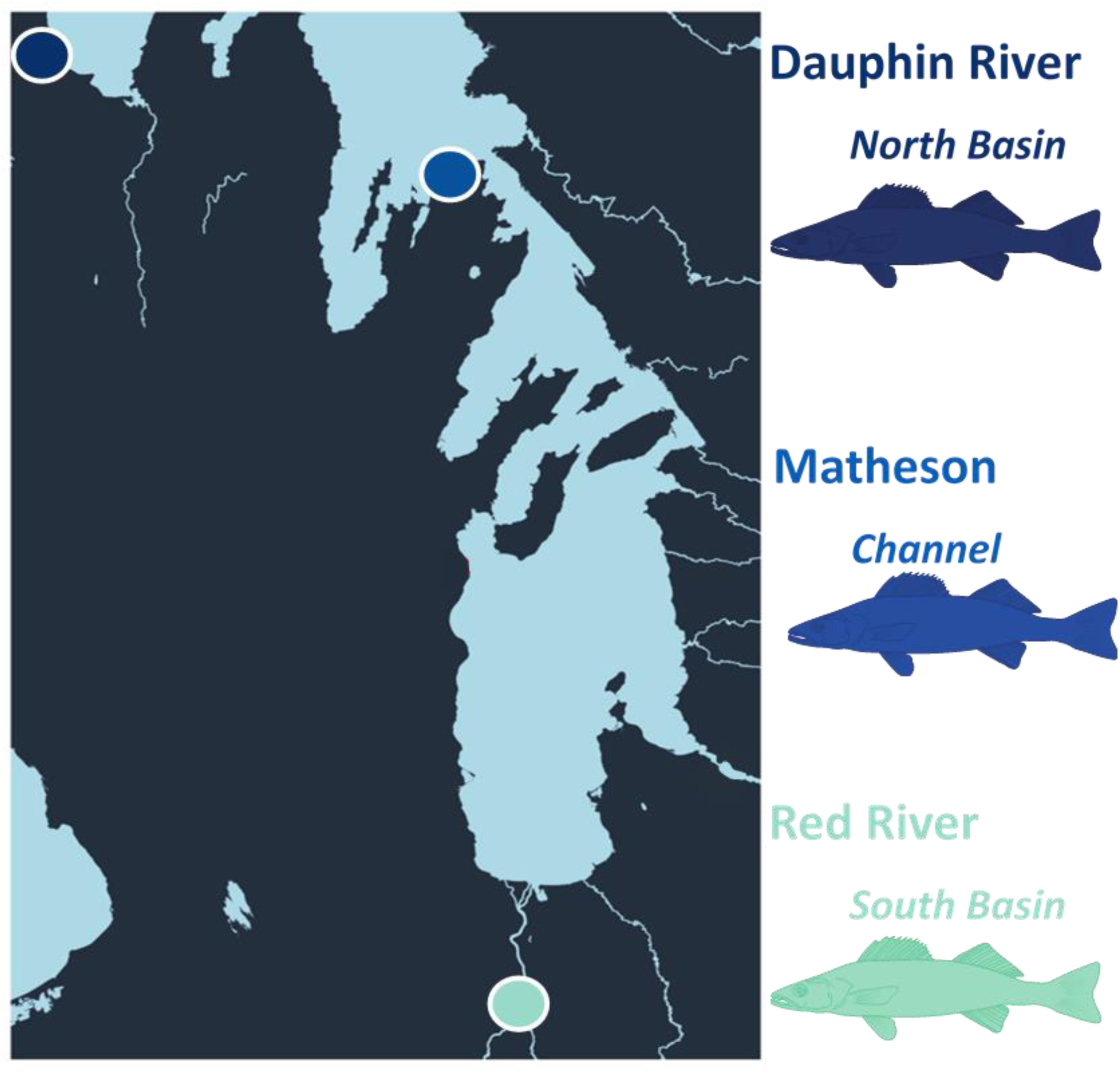
Sampling locations within Lake Winnipeg. Eight walleye (*Sander vitreus)* per year and per spawning site were collected, for *n* = 48 fish over 2017 and 2018. The Red River represents the south basin, Matheson Island represents the channel connecting the two lake basins, and the Dauphin River represents the north basin.

The current study aimed to show how mRNA sequencing can be an effective approach for developing critical pieces of information directly applicable to fisheries and conservation practitioners. We used RNA sequencing for genetic characterization of Lake Winnipeg walleye sampled from known spawning locations that potentially represent fish from the north and south basins. We also sampled fish collected at the channel that connects the north and south basins as an intermediate site. We hypothesized that walleye populations within Lake Winnipeg show evidence of distinct population differentiation identified using RNA sequencing data, despite the weak signatures from microsatellite data (Backhouse-James & Docker 2011). We predicted that the walleye population divergence may partially reflect the different environments and natural histories between the north and south basins of Lake Winnipeg.

## Methods

### RNA extraction and sequencing

Gill tissue was collected from large (≥ 1.2 kg) predominately female (44 female, 4 unidentified sex) (Supplementary Table 1) walleye over two years from three sites in the Lake Winnipeg system (Red River, Matheson Island, and Dauphin River, representing sites in the south basin, channel, and north basin, respectively; Figure 1; Supplementary Table 1; *n* = 8 per year and site, *n* = 48 total). These fish were sampled during the spawning season (approximately May through early June in 2017 and 2018). Walleye were collected by electrofishing, held in a live well for no longer than one hour, and anaesthetized using a Portable Electroanesthesia System (PES^™^, Smith Root, Vancouver, Washington, USA) in accordance with approved animal use protocols of Fisheries and Oceans Canada (FWI-ACC-2017-001, FWI-ACC-2018-001), the University of Manitoba (F2018-019) and the University of Nebraska-Lincoln (Project ID: 1208). Fish were sampled non-lethally for gill tissue, where 2–3 mm of the terminal ends of 3–4 filaments from the left side each fish were collected and placed in RNA*later* (Thermo Fisher Scientific, Waltham, Massachusetts, USA) that was kept at 4 °C for 24 h prior to storage at −80 °C. As part of a larger study on the physiological status and movement of Lake Winnipeg walleye, other samples collected were the first dorsal spine, scales, blood, a muscle biopsy, and fin clips. Fish were surgically implanted with VEMCO acoustic tags prior to release (VEMCO, Bedford, Nova Scotia, Canada). For the purposes of the present study, only gill tissue was analyzed. Total RNA extractions were performed on gill tissue using RNeasy Plus Mini Prep Kits (QIAGEN, Venlo, Netherlands) following manufacturer’s protocols, with minor modifications (provided in the supplementary materials).

The quantity and quality of RNA was assessed with a Nanodrop One Spectrophotometer (Thermo Fisher) and electrophoresis on a 1% agarose gel, respectively. Total RNA was normalized to 80 ng µL^-1^ and sent to the McGill University and Génome Québec Innovation Centre sequencing facility (http://gqinnovationcenter.com) for cDNA library preparation and sequencing. Total RNA was used to prepare 48 separate cDNA libraries to produce 100 base pair paired end reads using the NEBNext Ultra II Directional RNA Library Prep Kit for Illumina (New England Biolabs, Ipswich, Massachusetts, USA). Each library was individually barcoded with NEBNext dual adaptors (New England Biolabs) prior to sequencing. All 48 fish were sequenced on a single lane of a NovaSeq 6000 (Illumina, San Diego, California, USA). 2.17 billion reads total were sequenced, with an average of 45,225,548 reads per sample collected (5,071,090 s.d.) (Supplementary Table 1).

### SNP calling

Raw read files were uploaded to the Graham and Cedar clusters on the Westgrid section of the Compute Canada partnership (https://www.westgrid.ca/). Read files were checked for quality using FastQC version 0.11.8 (Andrews 2010) and trimmed using Trimmomatic version 0.36 (Bolger et al. 2014). When using FastQC version 0.11.8 (Andrews 2010), the program was set to allow two seed mismatches, a palindrome clip threshold of 30 nucleotides, and simple clip threshold of ten nucleotides. A sliding window size of 4 base pairs was used to filter data for a minimum Phred 64 quality of five, with five nucleotides trimmed from both the leading and trailing ends of reads, and a minimum read length of 36. After trimming, FastQC was used again to verify data quality. Scripts used for the analyses in this manuscript are provided at https://github.com/BioMatt/Walleye_RNAseq.

The SuperTranscripts pipeline was used to align reads (Davidson et al. 2017) for SNP calling. First, Salmon version 0.11.3 (Patro et al. 2017) was used to quantify read counts, as compared to a previously assembled reference transcriptome for walleye (Sequence Read Archive Accession SRP150633; Jeffrey et al, in revision). In Salmon, validate mappings, range factorization bins of size 4, sequencing bias, and GC bias options were all used, along with dumping equivalence classes for subsequent steps. Using the count estimates from Salmon, Corset version 1.07 (Davidson & Oshlack 2014) was used to cluster the data for assembly into SuperTranscripts. A linear representation of the transcriptome was constructed with Lace version 1.00 (https://github.com/Oshlack/Lace) using information from Corset and the original transcriptome, where 263,272 genes from the original transcriptome were gathered into 148,165 super clusters. Following Lace, STAR version 2.7.0a (Dobin et al. 2013) was used in 2-pass mode to align trimmed reads to the reassembled transcriptome. Here, annotated junctions from Lace were provided along with the new transcriptome, and sjdbOverhang of 99 was chosen following recommended settings of 1 base pair below read length. A minimum of 79.6% reads uniquely mapped to the Lace-clustered transcriptome (mean 81.5% ± 0.5% s.d.) (Supplementary Table 1).

For calling SNPs, the STAR-aligned reads were processed with Picard version 2.18.9 by adding read groups, splitting cigar ends, and merging bam files (Broad Institute 2018), then SNPs were called using FreeBayes version 1.2.0 (Garrison & Marth 2012). Detailed methods for calling SNPs are provided in the Supplementary Materials. We filtered the VCF file from FreeBayes in two ways. This resulted in 2,458,947 SNPs and 586,556 indels, which were used as unfiltered data for subsequent steps. To study SNPs as close to neutrality as possible, we used vcftools version 0.1.14 (Danecek et al. 2011) to filter for biallelic SNPs of genotype and site quality 30 with a minor allele frequency of 0.05 with no missing data, in Hardy-Weinberg Equilibrium with a *p*-value < 0.005. SNPRelate version 1.16.0 (Zheng et al. 2012) was then used to prune SNPs for linkage disequilibrium at a threshold of 0.20, where super clusters were coded as chromosomes for the purposes of linkage disequilibrium pruning. These steps resulted in a putatively neutral data set of 52,372 SNPs used for population structure analyses. For a broader subset of SNPs for which neutrality was not assumed, vcftools was used to filter for genotype and site with quality 30, minor allele frequency 0.05, and a maximum of two missing genotypes out of 48 possible. 222,634 SNPs were retained from these filtering steps, which were then used for outlier tests and functional analyses.

### Population structure

To investigate population structure using the 52,372 putatively neutral SNPs, we used a combination of exploratory analyses, either with no prior information or with sampling location provided as priors, and population reassignment and differentiation tests to find genetic clusters despite possible signals of admixture or gene flow. Structure version 2.3.4 (Falush et al. 2003; Falush et al. 2007; Hubisz et al. 2009; Pritchard et al. 2000) was run with no prior location or population information, an initial value of alpha of 1.0, a maximum value of alpha of 10.0, prior mean *F*_ST_ of 0.01, lambda of 1.0, a burn in period of 10,000 repetitions, and 110,000 Markov Chain Monte Carlo repetitions after burn in. Structure plots were visualized with pophelper version 2.2.7 (http://royfrancis.github.io/pophelper/). Ten replicates of K = 2–5 were tested.

For analyses performed in R (R Core Team 2019), the package vcfR was used to format genotype data for use with other programs (Knaus & Grünwald 2017). Adegenet version 2.1.1 (Jombart et al. 2010) was used in two ways. First, in an exploratory capacity to perform Discriminant Analysis of Principal Components (DAPC), where sampling location was provided for the DAPC as prior population information. Second, population structure was investigated irrespective of sampling location by using cluster identification from successive K-means, as implemented in the find.clusters function in Adegenet. Here, different numbers of clusters were explored in the data (40 principal components were retained for exploratory steps) and evaluated with a Bayesian Information Criterion (BIC), where the most well-supported number of clusters with lowest BIC was 2 (Supplementary Figure 1). In addition to exploring the two clusters, the population assignments from three clusters were used to explore genetic differentiation in the data because fish were sampled from three sites (Supplementary Table 1). With Hierfstat version 0.04-22 (Yang 1998; Weir & Cockerham 1984), the Weir & Cockerham’s pairwise *F*_ST_ was calculated among the three sampling locations, then between the two reassigned clusters described by Adegenet (Supplementary Table 1). We generated 95% confidence intervals for these *F*_ST_ values in Hierfstat using a bootstrap approach over 1,000 iterations.

To visualize population differentiation, we used a PCA as implemented in Adegenet version 2.1.1 (Jombart et al. 2010) and t-SNE as implemented in Rtsne version 0.15 (van der Maaten & Hinton 2008). For the t-SNE, two final dimensions were used, with 100 initial dimensions, 15 perplexity, theta of 0.5, and 5,000 iterations. These approaches were used with the same settings applied to the putatively neutral SNPs, and visualizations were thus comparable between data sets.

### Temporal stability & kinship

To test for temporal stability in the data, we created subsets of individuals caught in 2017 and 2018. As with the whole dataset, Weir & Cockerham’s pairwise *F*_ST_ was calculated both among sampling locations and between the two reassigned clusters, and generated 95% confidence intervals over 1,000 bootstrapped iterations in hierfstat (Supplementary Table 1). Modest results that are consistent over time support confidence in a real genetic signal, as opposed to results driven by bias which are more likely to be inconsistent over time (Waples 1998).

To address the possibility that sample collection, extraction, sequencing, or another process introduced an erroneous year effect into the data, we identified SNPs that differed between fish sampled in 2017 and 2018 with an *F*_ST_ above 0.01 using hierfstat, then filtered out those SNPs from the data using VCFtools version 0.1.14. Following these steps, 13,640 SNPs (26.04% of 52,372 neutral SNPs total) were identified as having a large effect between years and were thus removed, leaving 38,732 SNPs. Analyses for population structure were then re-run with this smaller set of SNPs. *F*_ST_ was calculated both between sites and between two reassigned clusters described by Adegenet (Supplementary Table 1). Data was also visualized by using a PCA as implemented in Adegenet version 2.1.1, and t-SNE as implemented in Rtsne version 0.15.

To test if our estimates of population structure were not driven by family groups (Waples 1998), we used Colony version 2.0.6.4 to reconstruct pedigrees in our sample of 48 individuals, with consideration of possible full-siblings (Jones & Wang 2010; Wang 2004). The putatively neutral SNPs were converted to the Colony format using a script by D. deWaters (https://github.com/dandewaters/VCF-File-Converter). The Colony command-line input file was then generated to run the program with updated allele frequencies, dioecy, inbreeding possible, polygamy allowed, no clones, full sibship scaling, no sibship prior, unknown population allele frequencies, ten runs of medium length, full likelihood inference, and high precision. This Colony input file was generated using a script originally written by M. Ackerman (used in Ackerman et al. 2017), modified and posted with permission for the present study. Independent of Colony’s maximum likelihood-based approach, we also used the method of moments as implemented in SNPRelate version 1.18.0 (Zheng et al. 2012) to estimate a kinship coefficient between individuals, also using the putatively neutral SNPs.

### Outlier SNPs

Using the full list of SNPs filtered for genotype quality 30, minor allele frequency > 0.05 and two missing individuals allowed, but not filtered for Hardy-Weinberg Equilibrium or Linkage Disequilibrium, we tested for outlier SNPs using an unsupervised approach in pcadapt version 4.1.0 (Luu et al. 2017). The unsupervised approach was used because weak population differentiation and the likely presence of admixed individuals in the data would either lower our sample size by filtering admixed individuals out, or lead to false-positive outlier loci by their inclusion when using a supervised approach with population structure included (Liu et al. 2016). While this may lead to issues of false positives from multiple tests (Foll & Gaggiotti 2008), we addressed this issue by using a q-value of 0.05 and focusing our interpretation on transcripts that contain two or more outlier SNPs. Two PCs were chosen for this analysis by observing the scree plot visualizing K = 1–20 following Cattell’s rule, where the point that a smooth decrease in eigenvalues levels off on a scree plot is the last important PC for explaining the data (Cattell 1966).

By relating the transcript ID of a significant outlier SNP (q-value < 0.05) to that transcript’s putative function and gene ID from the annotated reference transcriptome, a database of transcripts which diverged by sampling location or year was created for the Lake Winnipeg walleye in the present study. From this database, a list of transcripts relevant to either sampling location or year was used for gene set enrichment analysis using EnrichR (Chen et al. 2013; Kuleshov et al. 2016), thereby summarizing genes by gene ontology (GO) terms. In addition, transcripts were filtered to find those with two or more significant outlier SNPs that diverged by either sampling location or year, and these transcripts were few enough that enrichment analysis was not necessary. By only including genes with multiple outlier SNPs, we sought to reduce the presence of false positive signals in this outlier test.

## Results

### Population structure

Our data suggested weak but significant population structure between the north and south basins of Lake Winnipeg. The Red River and Matheson Island locations slightly diverged (*F*_ST_ = 0.0012), while the Dauphin River fish were the most genetically distinct group sampled (*F*_ST_ = 0.0068 and 0.0043 compared to the Red River and Matheson Island, respectively) (Figures 2, 3, Table 1, and Supplementary Figure 2). Moreover, Structure and the DAPC returned similar results with respect to which fish were admixed, although membership probabilities differed (Figure 2, 3). Between K = 2–5, Structure consistently separated the Dauphin River fish from the Matheson Island and Red River fish, while the Red River and Matheson Island fish did not separate from each other by site, but instead separated between years (Figure 2).

**Figure 2.**
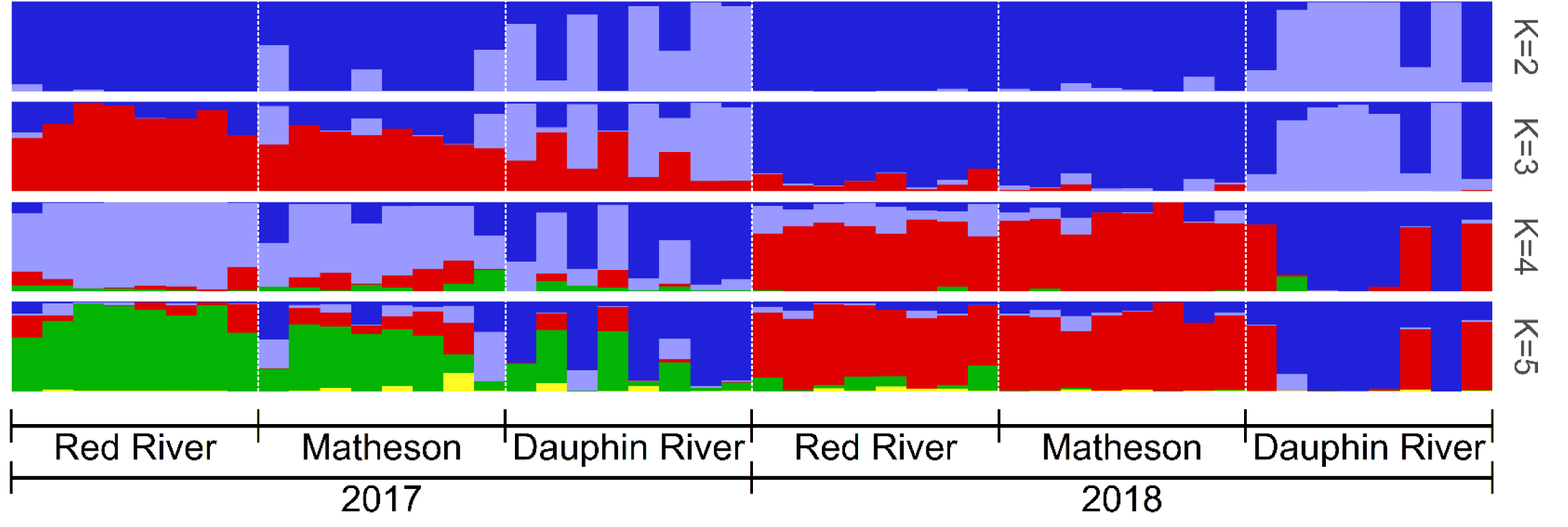
Representative Structure runs from ten replicates testing K = 2–5, organized by collection site (Red River in the south basin, Matheson Island in the channel, and Dauphin River in the north basin) and year collected (2017 and 2018) for all walleye (*Sander vitreus*) used in the present study. Collection site locations are available in Figure 1. This analysis was performed with 52,372 Hardy-Weinberg Equilibrium filtered and linkage disequilibrium pruned, putatively neutral single nucleotide polymorphisms.

**Table 1.**
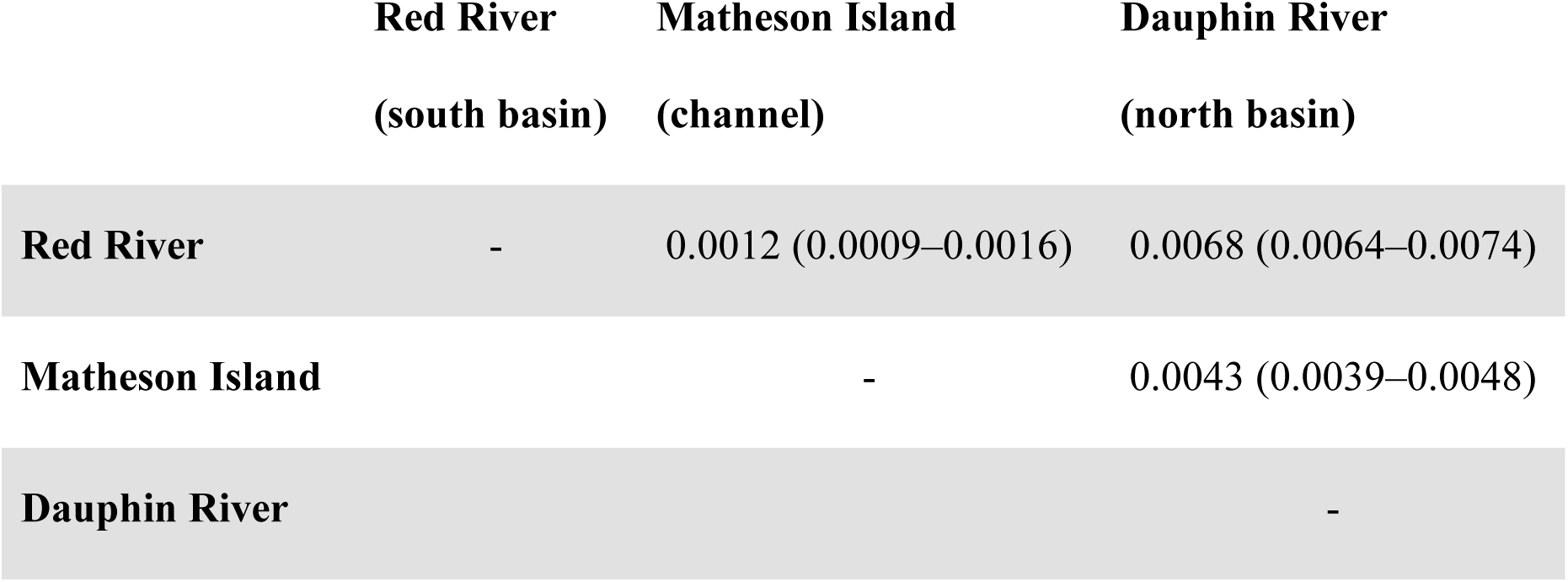
Weir & Cockerham’s pairwise *F*_ST_ calculated with hierfstat between the Red River in the south basin, Matheson Island in the channel, and Dauphin River in the north basin for all 48 walleye (*Sander vitreus*) sampled in both 2017 and 2018. 95% confidence intervals are provided in parentheses. Collection site locations are available in Figure 1. This analysis was performed with 52,372 Hardy-Weinberg Equilibrium filtered and linkage disequilibrium pruned, putatively neutral single nucleotide polymorphisms.

|  | **Red River**<br>(south basin) | **Matheson Island**<br>(channel) | **Dauphin River**<br>(north basin) |
| --- | --- | --- | --- |
| **Red River** | - | 0.0012 (0.0009–0.0016) | 0.0068 (0.0064–0.0074) |
| **Matheson Island** |  | - | 0.0043 (0.0039–0.0048) |
| **Dauphin River** |  |  | - |

The PCA and t-SNE used with the putatively neutral SNPs show similar patterns of Matheson and Red River fish separated, but more similar to each other than either with the Dauphin River fish (Supplementary Figure 2). When comparing the PCA and t-SNE plots between the neutral linkage disequilibrium-pruned SNPs and the broader collection of SNPs used for outlier analyses, genetic differentiation between the Red River and Matheson Island fish disappears when using all of the SNPs with the t-SNE, whereas separation between the two sites persists when only using Hardy-Weinberg Equilibrium filtered and LD-pruned SNPs.

### Population assignment

Using two clusters for reassignment from Adegenet, out of 48 fish, 36 clustered in one group (Cluster 1), and twelve in the other (Cluster 2; Supplementary Table 1). Cluster 1 was characterized by a combined Red River and Matheson Island group of fish with few Dauphin River fish (six were collected from the Dauphin River, 14 from Matheson Island, and 16 from the Red River), while Cluster 2 was characterized by Dauphin River fish and a small number of Matheson Island fish (ten fish from the Dauphin River and two from Matheson Island). Weir and Cockerham’s pairwise *F*_ST_ between these two reassigned clusters was 0.0095 with a 95% confidence interval between 0.0090–0.010.

Using three clusters for reassignments from Adegenet, out of 48 fish, 19 were in one group (Cluster 1), ten fish were in another group (Cluster 2), and 19 fish in a final group (Cluster 3). Clusters 1 and 3 were characterized by a year effect, where every individual in Cluster 1 was captured in 2018 and every individual in Cluster 3 was captured in 2017. Both Clusters 1 and 3 had 16 out of 19 fish coming from the Red River or Matheson Island sites. Meanwhile, all ten fish in Cluster 2 were from the Dauphin River with five fish each collected in 2017 and 2018 (Supplementary Table 1).

### Temporal stability and kinship

When partitioning individuals by sampling location and year collected, all confidence intervals for between-site pairwise *F*_ST_ estimates overlapped over both sampling years, indicating consistent patterns of between-site divergence in 2017 and 2018. However, values between the Dauphin River and Matheson Island varied the most, with an estimate of 0.0044 (0.0035–0.0052) in 2017, and 0.0060 (0.0051–0.0070) in 2018 (Table 2).

**Table 2.** Weir & Cockerham’s Pairwise *F*_ST_ calculated with hierfstat between the Red River in the south basin, Matheson Island in the channel, and Dauphin River in the north basin. *F*_ST_ values above the diagonal represent the 24 walleye (*Sander vitreus*) collected in 2017, and values below the diagonal represent the 24 collected in 2018. 95% confidence intervals are provided in parentheses. Collection site locations are available in Figure 1. This analysis was performed with 52,372 Hardy-Weinberg Equilibrium filtered and linkage disequilibrium pruned, putatively neutral single nucleotide polymorphisms.

|  | Red River<br>(south basin) | Matheson Island<br>(channel) | Dauphin River<br>(north basin) |
| --- | --- | --- | --- |
| Red River | - | 0.0023 (0.0015–0.0031) | 0.0073 (0.0064–0.0082) |
| Matheson Island | 0.0019 (0.0011–0.0028) | - | 0.0044 (0.0035–0.0052) |
| Dauphin River | 0.0067 (0.0058–0.0077) | 0.0060 (0.0051–0.0070) | - |

Using the 38,732 SNPs filtered for loci, which showed *F*_ST_ between years of > 0.01, *F*_ST_ between the two reassigned clusters found using Adegenet (Supplementary Table 1) was 0.010 (0.0094–0.011). With these same year effect-filtered SNPs, pairwise *F*_ST_ between sites did not significantly differ from values found using the neutral SNPs either overall or in a subset by year (Supplementary Table 2). The PCA and t-SNE on the SNPs filtered for a year effect showed patterns of spatial differentiation consistent with other analyses, with the Dauphin River fish being more separate from the Red River and Matheson Island group of fish (Supplementary Figure 3).

We found no evidence of kinship using either Colony or the method of moments. Over ten replicate runs in Colony, individuals belonged to separate families with inclusive and exclusive probabilities of 1.0000 each. Using the method of moments implemented in SNPRelate (Zheng et al. 2012), the highest kinship coefficient between two individuals was 0.096 (mean 0.053 ± 0.019 s.d.), where a kinship coefficient of approximately 0.5 would indicate full-siblings.

### Outlier SNPs

There was site-specific differentiation across Principal Component 1 (PC1) in the pcadapt analysis (Figure 4). In total, 1,177 SNPs were outliers at q < 0.05, with 386 SNPs contributing to PC1 where fish separated by site, and 791 SNPs contributing to PC2 where fish separated by year (Figure 4). For the 386 SNPs associated with PC1 (Figure 4), 120 uniquely annotated transcripts were available for enrichment analysis using EnrichR. These transcripts corresponded to GO terms such as purine ribonucleoside triphosphate binding, ATP binding, and adenyl ribonucleotide binding, all significant at Benjamini-Hochberg adjusted *p*-values < 0.05 (Supplementary Table 3). By filtering for uniquely annotated transcripts with ≥ 2 outlier SNPs associated with PC1, 19 transcripts were identified (Table 3) that varied by sampling location. Six of these genes were associated with ion channels and cell membrane transport, including claudin-10, ankyrin-3, sodium/hydrogen exchanger 6, sodium/potassium-transporting ATPase subunit alpha-3, perforin-1, and ATP-binding cassette sub-family A member 12. Additionally, four genes that varied spatially were associated with the cytoskeleton, such as myosin-9, beta/gamma crystallin domain-containing protein 1, tubulin beta-4B chain, and interferon-induced protein 44.

**Figure 3.**
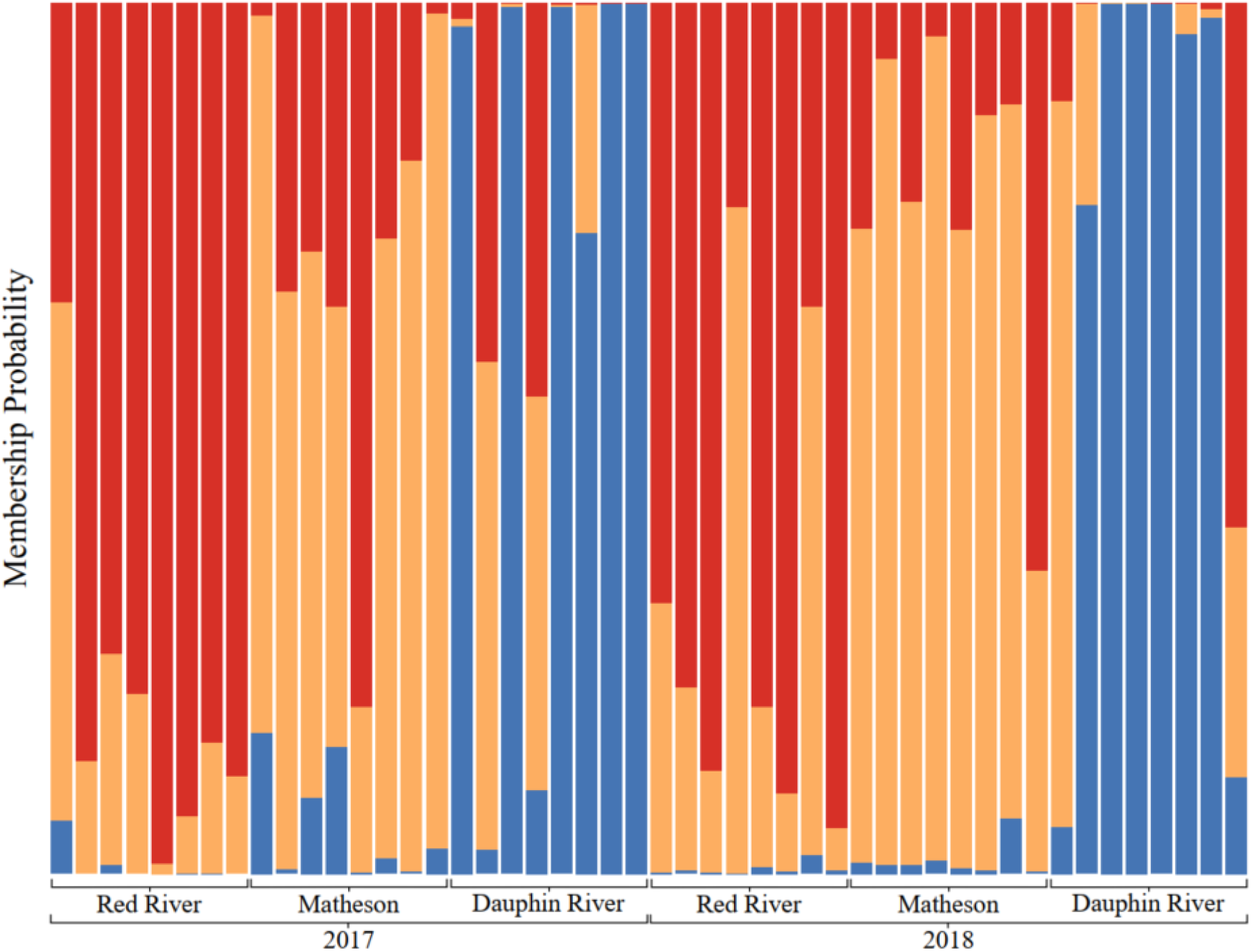
Membership probability plot of discriminant analysis of principal components using prior collection site information (Red River in the south basin, Matheson Island in the channel, and Dauphin River in the north basin) on walleye (*Sander vitreus*) collected over 2017 and 2018, performed using Adegenet. Collection site locations are available in Figure 1. This analysis was performed with 52,372 Hardy-Weinberg Equilibrium filtered and linkage disequilibrium pruned, putatively neutral single nucleotide polymorphisms.

**Figure 4.**
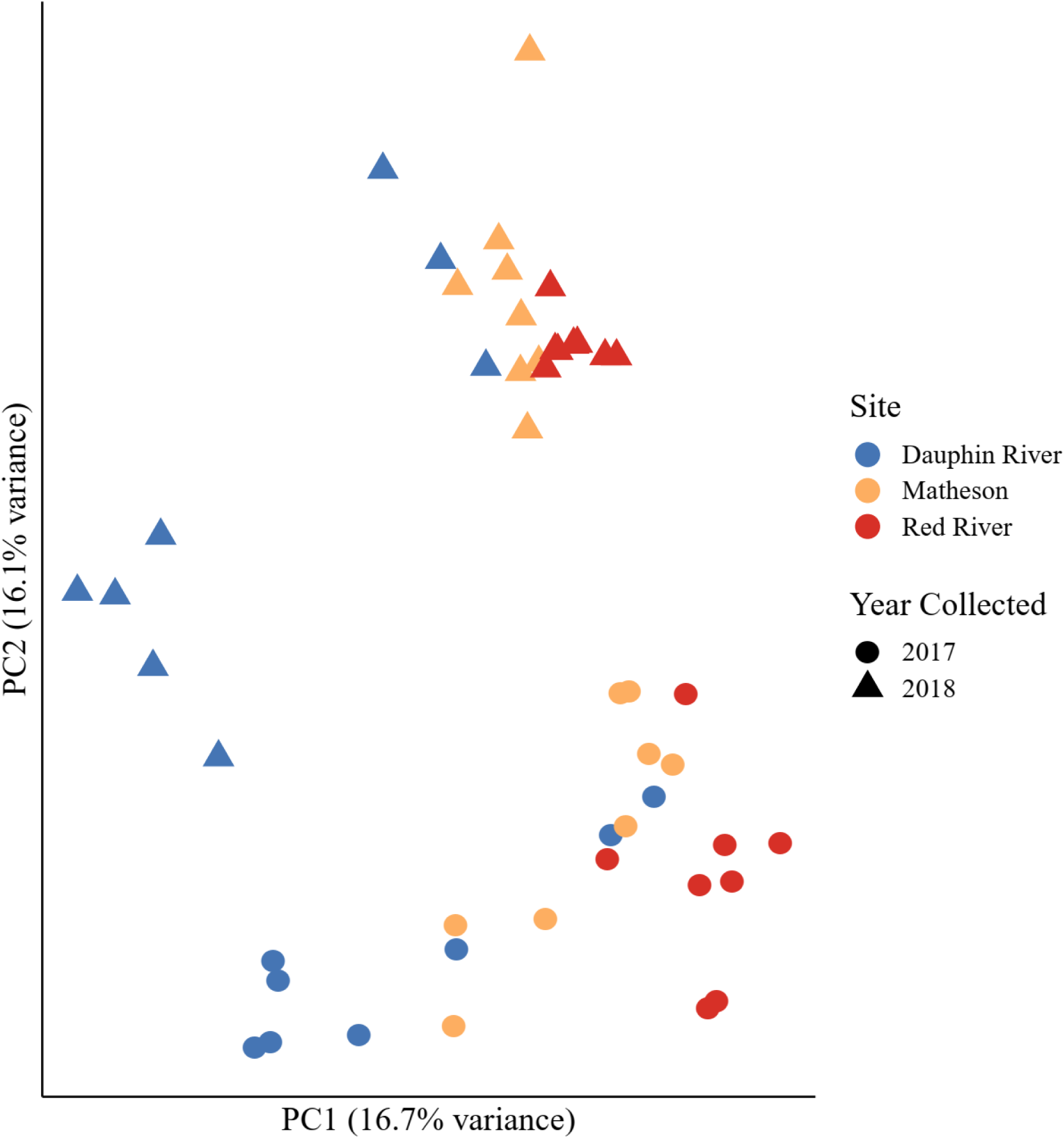
Principal Components Analysis implemented in pcadapt with color showing site collected (red for Red River in the south basin, yellow Matheson Island in the channel, and blue Dauphin River in the north basin), circles showing walleye (*Sander vitreus*) collected in 2017, and triangles showing walleye collected in 2018. Collection site locations are available in Figure 1. This analysis was performed using a set of 222,634 single nucleotide polymorphisms (SNPs) filtered for quality, minor allele frequency > 0.05, and a maximum of two out of 48 missing individuals, but not filtered for Hardy-Weinberg Equilibrium or pruned for linkage disequilibrium. These SNPs are, thus, more likely to represent patterns of adaptive variation in the system, and outlier analyses were performed using this set of SNPs. Principal Component 1 (PC1) represents a latitudinal gradient, while Principal Component 2 (PC2) represents a genetic divergence between sampling years. Outlier SNPs that contribute to each of these axes were selected for functional analyses (Tables 3, 4, 5), with 386 SNPs contributing to PC1 and 791 SNPs contributing to PC2, significant at Benjamini-Hochberg adjusted *p*-values < 0.05.

**Table 3.**
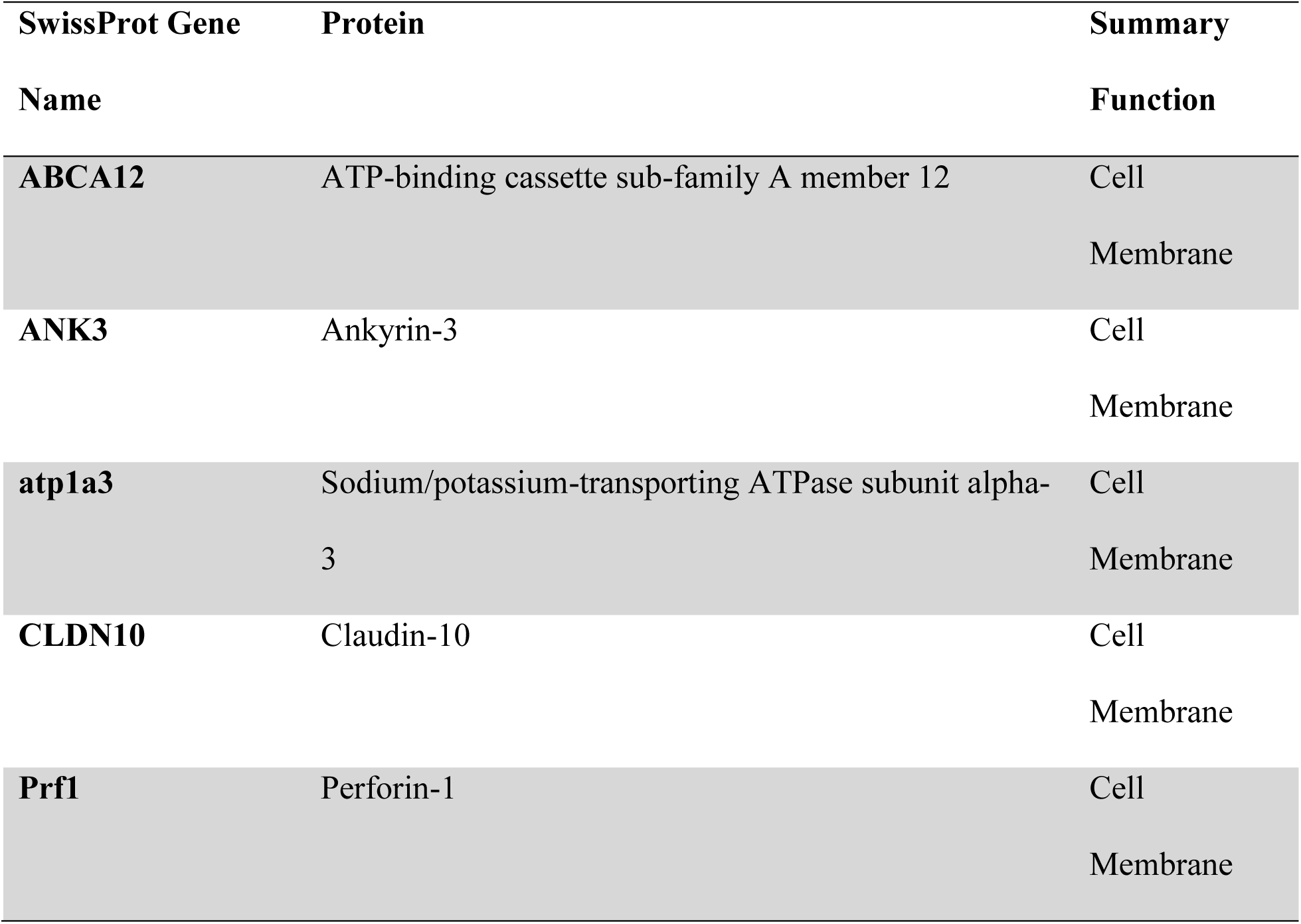
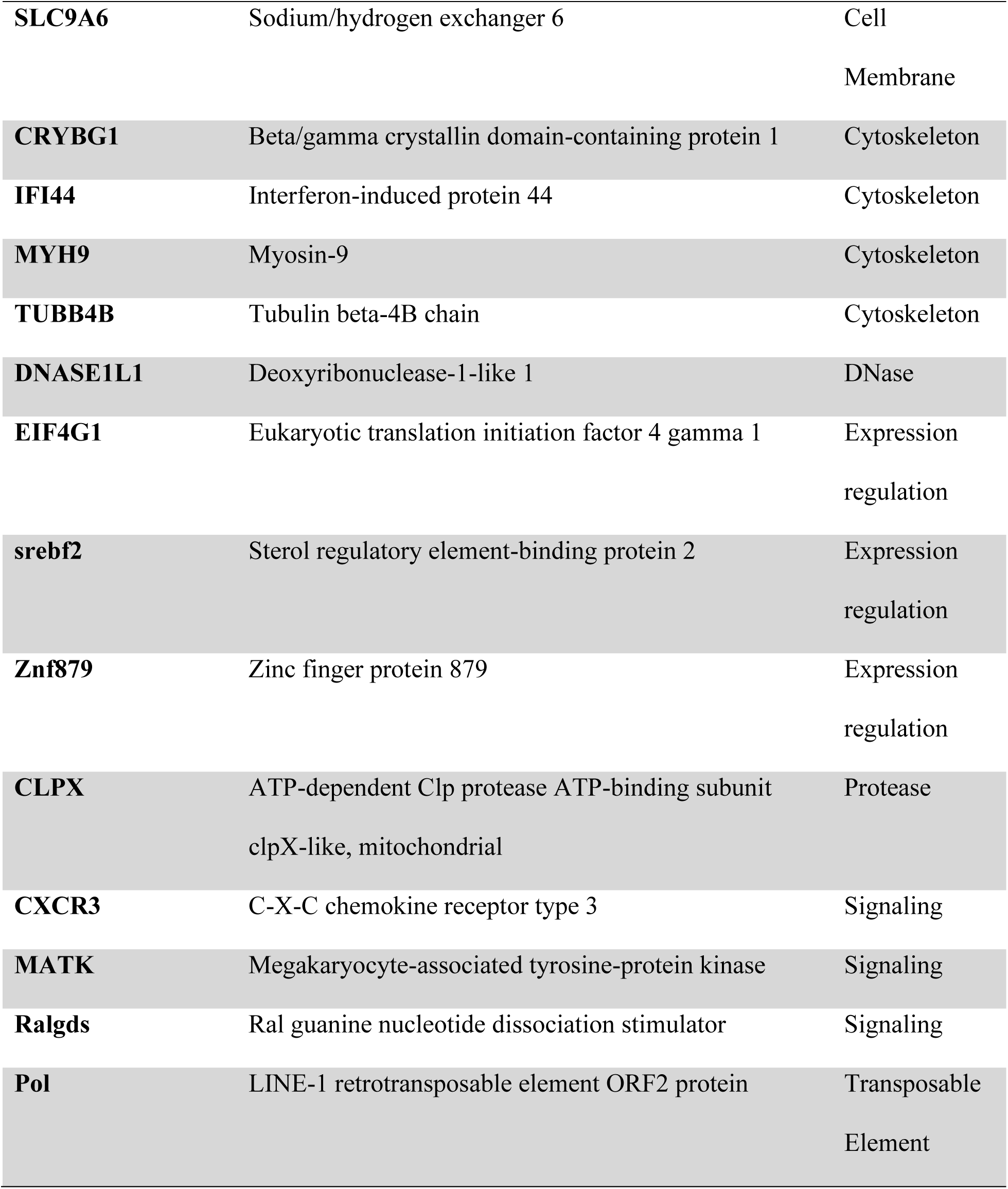
Genes that vary along a latitudinal gradient in Lake Winnipeg walleye (*Sander vitreus*) with ≥ 2 outlier single nucleotide polymorphisms (SNPs) from pcadapt, each significant at a Benjamini-Hochberg adjusted *p*-value < 0.05 (PC1 in Figure 4). SwissProt gene names and corresponding proteins are provided, and general cellular location or function of these genes are described in Summary Function. This analysis was performed using a set of 222,634 single nucleotide polymorphisms (SNPs) not filtered for Hardy-Weinberg Equilibrium or pruned for linkage disequilibrium, unlike the putatively neutral set of SNPs used for population structure analyses.

**Table 4.** Genes that vary between 2017 and 2018 in Lake Winnipeg walleye (*Sander vitreus*) with transcripts containing ≥2 outlier single nucleotide polymorphisms (SNPs) from pcadapt, each SNP significant at a Benjamini-Hochberg adjusted *p*-value < 0.05 (PC2 in Figure 4). SwissProt gene names and corresponding proteins are provided, and general cellular location or function of these genes are described in Summary Function. This analysis was performed using a set of 222,634 single nucleotide polymorphisms (SNPs) not filtered for Hardy-Weinberg Equilibrium or pruned for linkage disequilibrium, unlike the putatively neutral set of SNPs used for population structure analyses.

| SwissProt Gene<br>Name | Protein | Summary Function |
| --- | --- | --- |
| **Gp1bb** | Platelet glycoprotein Ib beta chain | Cell Adhesion |
| **MFAP4** | Microfibril-associated glycoprotein 4 | Cell Adhesion |
| **BTG3** | Protein BTG3 | Cell Division |
| **Ppp6c** | Serine/threonine-protein phosphatase 6 catalytic subunit | Cell Division |
| **Slc12a3** | Solute carrier family 12 member 3 | Cell Membrane |
| **MMD** | Monocyte to macrophage differentiation factor | Expression regulation |
| **Srsf10** | Serine/arginine-rich splicing factor 10 | Expression regulation |
| **Ube2a** | Ubiquitin-conjugating enzyme E2 A | Expression regulation |
| **Znf18** | Zinc finger protein 18 | Expression regulation |
| **St6galnac2** | Alpha-N-acetylgalactosaminide alpha-2,6-sialyltransferase 2 | Protein Modification |
| **MYO9A** | Unconventional myosin-IXa | Signaling/Cytoskeleton |
| **Pol** | LINE-1 retrotransposable element ORF2 protein | Transposable Element |
| **RTase** | Probable RNA-directed DNA polymerase from transposon BS | Transposable Element |
| **TC1A** | Transposable element Tc1 transposase | Transposable Element |
| **TN6** | Putative transposase in *Dicentrarchus labrax* (European seabass) | Transposable Element |
| **TY3B-G** | Transposon Ty3-G Gag-Pol polyprotein | Transposable Element |
| **YTX2** | Transposon TX1 uncharacterized protein (Fragment) | Transposable Element |

Using the 791 SNPs associated with Principal Component 2 (PC2), which varied by year (Figure 4), 130 uniquely annotated transcripts were available for enrichment analysis; however, no GO terms were significant at an adjusted *p*-value < 0.05. For transcripts with ≥ 2 PC2 outlier SNPs, 17 uniquely annotated genes were identified of which six were either transposons, transposable elements, or fragments of transposons (Table 4). Two genes that code for the proteins serine/threonine-protein phosphatase 6 catalytic subunit and protein BTG3, which regulate cell division in the G1 to S phase transition were also identified (Table 4).

## Discussion

We observed weak population structure characterized by groups collected at the Red River and Matheson Island sampling locations, representing south basin and channel fish, contrasted with a group collected at the Dauphin River, representing north basin fish. As such, the north and south basin walleye in Lake Winnipeg may be separate groups with an *F*_ST_ of 0.0095, but with gene flow between them primarily at the channel connecting the two basins. Consistent with results in the present study, a study using microsatellites found a similar weak, but significant, differentiation between Lake Winnipeg walleye from sites in the north and south basins (e.g., *F*_ST_ = 0.022 between the Grand Rapids in the north and Red River in the south; see Backhouse-James & Docker 2012), suggesting genomic divergence between walleye from the two basins. One important factor that may have contributed to weak population structure are historical stocking programs, which may have introduced walleye from nearby Lake Manitoba to Lake Winnipeg (Backhouse-James & Docker 2012). This unknown amount of gene flow from other systems, including up to 26.5 million fish annually between 1970 and 1983 (Lysack 1986), may have masked signatures of spatial population differentiation in Lake Winnipeg.

### Temporal differentiation

Differentiation between years was strongest in the south basin, where the Red River and Matheson Island fish separated by year to a greater extent than the Dauphin River walleye. Three hypotheses may explain these patterns of stronger temporal differentiation in the south basin. First, a cohort effect may underlie this pattern, where different year classes were more strongly represented in the lake during a given sampling year. A cohort effect could be the result of greater fishing pressure in the south basin than the north basin, as indicated by smaller allowed net mesh sizes in the south basin (Manitoba Sustainable Development 2019). Fishing pressure can change population dynamics and age structure in exploited species (Anderson et al. 2008; Murphy et al. 2001), therefore large fisheries operating since at least 1890 may have affected age structure in Lake Winnipeg fish (Department of Fisheries 1891). Cohort effects may alternatively be influenced by environmental conditions including predation intensity, water temperature, and time to hatch as observed in Lake Erie, Oneida Lake, and Lake Huron walleye (Busch et al. 1975; Fielder et al. 2007; Forney 1976). A second hypothesis is that some Lake Winnipeg walleye may engage in unobserved skipped spawning or alternate year spawning, which have been unexpectedly found in several species and may be present in walleye (Carlander et al. 1960; Henderson et al. 1996; Moles et al. 2008; Rideout & Tomkiewicz 2011). Third, the observed year effect may be an artifact of error introduced during sampling, extraction, sequencing, or bioinformatics. While some error contributing to between-year differentiation is impossible to rule out, the possibility that a particular analysis or filtering method introduced the year effect is reasonably small given that distinct tests showed consistent year effects. Moreover, the data reveal a consistency in spatial patterns with and without the year effect, demonstrating that at least the spatial population differentiation in Lake Winnipeg walleye is likely real.

While pairwise site *F*_ST_ values were temporally consistent (i.e., no significant differences between years), the greatest pairwise *F*_ST_ confidence interval difference between years was between walleye collected at Matheson Island and the Dauphin River, where confidence intervals for *F*_ST_ estimates overlapped by only 0.0001 between 2017 and 2018. Following Amrhein et al. (2019), we interpret here the possibility that the entire range of these confidence intervals reflect meaningful patterns in the data, and that *F*_ST_ between the Dauphin River and Matheson Island was different between 2017 and 2018. Because Matheson Island represents a narrow channel connecting the north and south basins of Lake Winnipeg, fish which would normally spawn in the Dauphin River may have used the channel more often in 2017, thus, lowering *F*_ST_ when performing a site-wise comparison. This difference in habitat use may have arisen from an undetermined environmental variable, such as time of ice melt, which in the north basin was ten days later in 2018 than in 2017 (D. Watkinson, unpublished data found using https://zoom.earth/).

Notably, gene flow appears to be one way from the southern Red River, northward. Going by capture location, no fish caught in the Red River showed a genetic background consistent with the Dauphin River fish, while with the Adegenet-reassigned clusters, no fish assigned to the mostly Dauphin River group was found in the Red River. On the other hand, fish which showed a genetic background consistent with the Red River group were found in the Dauphin River, both based on capture location and population reassignment.

### Biological significance

Several studies report morphological and life history differences between basins in Lake Winnipeg walleye consistent with the two delineated groups found in this study. Furthermore, environmental data show a north-south basin distinction with temperature, turbidity, mean depth, suspended solid, sulphate, sodium, chloride, and nutrient differences between the two basins (Brunskill et al. 1980; Environment Canada 2011). Walleye in the south basin show a bimodal growth pattern, where fisheries-induced selection may have contributed to the observation of dwarf walleye (Johnston et al. 2012; Mole et al. 2010; Sheppard et al. 2018). Harvest-induced genetic changes have been linked to size reductions in other walleye within two generations (Bowles et al. 2019). If walleye were panmictic throughout Lake Winnipeg, we might expect the dwarf morphotype to occur with similar frequency in the north basin. However, out of 616 total walleye caught in 2010 and 2011 (178 in the north basin, 438 in the south basin), only two out of 32 dwarf fish were caught in the north basin (Sheppard et al. 2018). Diet has also been shown to differ between north and south basin walleye, possibly because of prey or turbidity differences between the two basins (i.e., higher turbidity in the south basin) (Brunskill et al. 1980; Sheppard et al. 2015). Between 1979 and 2003, population characteristics such as age and length at 50% maturity were higher, while growth rate was slower in the north basin walleye, suggesting some level of isolation among walleye between basins (Johnston et al. 2012). These population characteristics may no longer be higher in the north basin following the collapse of the rainbow smelt (*Osmerus mordax)*, after which walleye body condition has decreased since 2010 (Manitoba Government 2018). Scale morphometry further suggests differences among spawning aggregations of walleye, especially between the north and south basins (Kritzer & Sale 2004; Watkinson & Gillis 2005). Taken together, the results of our study and those of previous studies suggest weak population structure among Lake Winnipeg walleye, with differentiation between walleye in the north and south basins. This pattern of weak population structure, high connectivity, but biologically significant differentiation is common in marine fishes such as the Atlantic cod (*Gladus morhua*) or Atlantic salmon (*Salmo salar*) (Aykanat et al. 2015; Knutsen et al. 2011), and of other walleye such as those observed in Lake Erie (Chen et al. 2019; Stepien et al. 2018).

The results of the present study suggest that the genetic differences between Lake Winnipeg walleye populations may have functional consequences. Out of 19 transcripts that had multiple SNPs that varied by sampling location, eight were related to membrane function, particularly ion channel activity. One of these proteins, Claudin-10 mRNA expression levels have been related ammonia exposure (Connon et al. 2011), rearing density (Sveen et al. 2016), and salinity (Bossus et al. 2015; Kolosov et al. 2013; Marshall et al. 2018) in fishes. Spatial variation in cell membrane proteins is consistent with environmental differences between basins in chemicals such as sodium, chloride, and phosphorous (Environment Canada 2011), although the biological impacts of these spatial chemical differences is unknown. Four cytoskeletal proteins were represented in the outlier SNPs that vary by sampling location as well. Cytoskeletal function is connected to cell growth in plants (Hussey et al. 2006; Wasteneys & Galway 2003), yeast (Li et al. 1995; Pruyne & Bretscher 2000), mouse cells (Kim et al. 2006; Kim & Coulombe 2010), and zebrafish (Johnston et al. 2011). Spatial variation in genes related to cell growth may thus be consistent with growth rate differences observed among walleye in Lake Winnipeg, where north basin fish had higher growth rates in 2010 and 2011 (Sheppard et al. 2018).

### Limitations

Despite its advantages, there are some limitations to using mRNA sequencing in the context of population genetics. The depth of sequencing required for differential gene expression and differential exon usage leads to greater costs associated with mRNA sequencing studies relative to reduced representation methods such as RAD-seq (Davey & Blaxter 2011). This often translates to a lower sample size, as is the case in our study. Reduced sample sizes can bias aspects of population genetic analyses, including identifying population structure (Waples & Gaggiotti 2006) and outlier SNPs (Luu et al. 2016). Second, mutations in mRNA are widely under selection (Chamary & Hurst 2005), therefore, caution must be exercised when interpreting SNPs from mRNA in genetic tests assuming neutrality. Third, linkage disequilibrium is useful for analyses of selective sweeps and demographic history (Catchen et al. 2017; Garrigan & Hammer 2006; Hoffmann & Willi 2008), among other approaches, but mRNA data may not be appropriate for these analyses because the extent of linkage between transcripts and marker density across the genome may be unknown. Finally, one key element of how transcriptomics was used in the present study is that it measured expressed mRNA in gill tissue. Messenger RNA expression provides useful information for transcript quantification-based analyses, but likely biases SNP discovery toward more highly expressed transcripts in the tissue collected. It is unknown how this expression-specific bias may influence population genomics. Nevertheless, mRNA sequencing has proven useful for recapitulating population structure discovered with traditional genetic methods (Jeffries et al. 2019) and describing previously uncharacterized population structure (Ellison et al. 2011; Yan et al. 2017).

### Conservation applications

We used population genetics and outlier detection to characterize weak, but biologically significant population structure, possible one-way gene flow, and genetic variation possibly underlying biological differences among Lake Winnipeg walleye. These results are consistent with observations of behavioural differences leading to fine-scale divergence in the walleye of other systems (Stepien et al. 2009). The low levels of population differentiation and possible gene flow from the south basin northward, indicate that this system may be effectively managed as a demographically connected metapopulation with two contributing populations (Kritzer & Sale 2004), consistent with conclusions from scale morphology presented in Watkinson & Gillis (2005) and with observations of subtle stock structure in Lake Erie walleye (Chen et al. 2019; Stepien et al. 2018).

The results from this study provide valuable information for walleye management, especially because the status of Lake Winnipeg walleye is becoming a concern and conservation action may be necessary to sustain the fishery. Signs of a declining fishery include a decrease in biomass and body condition between 2010 and 2015 (Manitoba Government 2018), possible unnatural selection against larger, economically desirable fish (Allendorf & Hard 2009; Bowles et al. 2019; Moles et al. 2010), models showing walleye harvests have been above maximum sustainable yields since the early 2000s, and a trend in harvest decline since 2010 (Manitoba Sustainable Development 2017–2018). The data gathered here, particularly the spatial variation in genes that may drive functional differences among Lake Winnipeg walleye, is useful for generating hypotheses that test and explain organismal responses to environmental stressors, thereby providing additional information for resource managers. For instance, life-history trait differences can inform conservation in threatened fishes by identifying resilient populations in a system (Hamidan & Britton 2015). Therefore, possible functional variation identified in this study may underlie heritable genetic differences among Lake Winnipeg walleye that change important traits such as tolerance to environmental conditions and growth rate differences. This information may be useful for integrating demographic connectivity and functional differences among walleye into a cohesive management framework.

We have shown how RNA sequencing data can be used for a population genomic scan in a non-model fish, even in a system where little molecular information is available. Filtering for Hardy-Weinberg equilibrium and linkage disequilibrium allows investigators to draw neutral markers from mRNA sequence data, making it useful for classical population genetic approaches. By contrast, the wide selective effects present in species’ transcriptomes allow for hypothesis-generating outlier tests that may reveal variation underlying phenotypic differences among populations. Non-lethal sampling makes RNA sequencing useful for species with low population sizes and for follow-up studies, such as the potential to track tagged individuals from which tissue has been collected. Because RNA sequencing data can yield physiological in addition to genetic information discussed here, we argue that it is useful for addressing diverse molecular questions in the conservation of freshwater species.

## Supporting information

All Supplementary Materials

## Acknowledgements

We thank D. Watkinson, C. Charles, C. Kovachik, D. Leroux, N. Turner, M. Gaudry, S. Glowa, and E. Barker for helping to sample these fish in the field. D. Watkinson and C. Charles also assisted with ice on/ice off data and a map of Lake Winnipeg, respectively. Dr. C. Garroway provided valuable suggestions in data analysis, and Dr. G. Anderson engaged in useful conversations about Lake Winnipeg ecology. We thank the Lake Winnipeg Research Consortium Inc. for supporting the open access publication of this manuscript. Many analyses were enabled by our chance to use computing resources provided by WestGrid (www.westgrid.ca) and Compute Canada (www.computecanada.ca). Sequencing was performed at the McGill University and Génome Québec Innovation Centre sequencing facility. This work was supported by a Fisheries and Oceans Canada Ocean and Freshwater Science Contribution Program Partnership Fund grant awarded to J.R.T., K.M.J. and Darren Gillis, and Natural Sciences and Engineering Research Council of Canada Discovery Grants awarded to K.M.J. (#05479) and J.R.T. (#06052). Work by J.R.T. is also supported by the Canada Research Chairs program (#223744) and the Faculty of Science, University of Manitoba (#319254).

## Data Archiving Statement

Raw sequence reads are available through the National Center for Biotechnology Information Sequence Read Archive (accession #PRJNA596986, https://www.ncbi.nlm.nih.gov/sra/PRJNA596986).

## Works Cited

Ackerman, M. W., Hand, B. K., Waples, R. K., Luikart, G., Waples, R. S., Steele, C. A., Garner, B.A., McCane, J., Campbell, M. R. (2017). Effective number of breeders from sibship reconstruction: empirical evaluations using hatchery steelhead. Evolutionary Applications, 10(2), 146–160. https://doi.org/10.1111/eva.12433

Amrhein, V., Greenland, S., & McShane, B. (2019). Scientists rise up against statistical significance. Nature, 567(7748), 305–307. https://doi.org/10.1038/d41586-019-00857-9

Anderson, C. N. K., Hsieh, C. H., Sandin, S. A., Hewitt, R., Hollowed, A., Beddington, J., May, R., Sugihara, G. (2008). Why fishing magnifies fluctuations in fish abundance. Nature, 452(7189), 835–839. https://doi.org/10.1038/nature06851

Andrews, S. (2010). FastQC: a quality control tool for high throughput sequence data. https://www.bioinformatics.babraham.ac.uk/projects/fastqc/

Aykanat, T., Johnston, S. E., Orell, P., Niemelä, E., Erkinaro, J., & Primmer, C. R. (2015). Low but significant genetic differentiation underlies biologically meaningful phenotypic divergence in a large Atlantic salmon population. Molecular Ecology, 24(20), 5158–5174. https://doi.org/10.1111/mec.13383

Backhouse-James, S. M., & Docker, M. F. (2012). Microsatellite and mitochondrial DNA markers show no evidence of population structure in walleye (*Sander vitreus*) in Lake Winnipeg. Journal of Great Lakes Research, 38, 47–57. https://doi.org/10.1016/j.jglr.2011.05.005

Bolger, A. M., Lohse, M., & Usadel, B. (2014). Trimmomatic: A flexible trimmer for Illumina sequence data. Bioinformatics, 30(15), 2114–2120. https://doi.org/10.1093/bioinformatics/btu170

Bowles, E., Marin, K., & Fraser, D. J. (2019). Size reductions and genomic changes associated with harvesting within two generations in wild walleye populations. BioRxiv Preprint, (September). https://doi.org/10.1101/787374

Braje, T. J., & Erlandson, J. M. (2013). Human acceleration of animal and plant extinctions: A Late Pleistocene, Holocene, and Anthropocene continuum. Anthropocene, 4, 14–23. https://doi.org/10.1016/j.ancene.2013.08.003

Broad Institute. (2019). Picard Tools. Broad Institute, GitHub Repository. Retrieved from http://broadinstitute.github.io/picard/

Brunskill, G.J., Elliot, S.E.M., & Campbell, P. (1980). Morphometry, hydrology, and watershed data pertinent to the limnology of Lake Winnipeg. Canadian Manuscript Report of Fisheries & Aquatic Sciences No. 1556. 1–32.

Busch, W.-D. N., Scholl, R. L., & Hartman, W. L. (1975). Environmental factors affecting the strength of walleye (*Stizostedion vitreum vitreum*) year-classes in western Lake Erie, 1960–70. Journal of the Fisheries Research Board of Canada, 32(10), 1733–1743. https://doi.org/10.1139/f75-207

Carlander, K. D., Whitney, R. R., Speaker, E. B., & Madden, K. (1960). Evaluation of Walleye Fry Stocking in Clear Lake, Iowa, by Alternate-Year Planting. Transactions of the American Fisheries Society, 89(3), 249–254. https://doi.org/10.1577/1548-8659(1960)89[249:EOWFSI]2.0.CO;2

Catchen, J. M., Hohenlohe, P. A., Bernatchez, L., Funk, W. C., Andrews, K. R., & Allendorf, F. W. (2017). Unbroken: RADseq remains a powerful tool for understanding the genetics of adaptation in natural populations. Molecular Ecology Resources, 17(3), 362–365. https://doi.org/10.1111/1755-0998.12669

Chamary, J., & Hurst, L. D. (2005). Evidence for selection on synonymous mutatations affecting stability of mRNA secondary structure in mammals. Genome Biology, 6(9), R75. https://doi.org/10.1186/gb-2005-6-9-r75

Chen, K. Y., Euclide, P. T., Ludsin, S. A., Larson, W., Sovic, M. G., Gibbs, H. L., & Marschall, E. A. (2019). RAD-seq refines previous estimates of genetic structure in Lake Erie walleye (*Sander vitreus*). Transactions of the American Fisheries Society. https://doi.org/10.1002/tafs.10215

Chen, E. Y., Tan, C. M., Kou, Y., Duan, Q., Wang, Z., Meirelles, G., … Ma’ayan, A. (2013). Enrichr: Interactive and collaborative HTML5 gene list enrichment analysis tool. BMC Bioinformatics, 14(1), 128. https://doi.org/10.1186/1471-2105-14-128

Cingolani, P., Platts, A., Wang, L. L., Coon, M., Nguyen, T., Wang, L., … Ruden, D. M. (2012). A program for annotating and predicting the effects of single nucleotide polymorphisms, SnpEff. Fly, 6(2), 80–92. https://doi.org/10.4161/fly.19695

Connon, R. E., Jeffries, K. M., Komoroske, L. M., Todgham, A. E., & Fangue, N. A. (2018). The utility of transcriptomics in fish conservation. The Journal of Experimental Biology, 221(2), jeb148833. https://doi.org/10.1242/jeb.148833

Danecek, P., Auton, A., Abecasis, G., Albers, C. A., Banks, E., DePristo, M. A., … 1000 Genomes Project Analysis Group. (2011). The variant call format and VCFtools. Bioinformatics, 27(15), 2156–2158. https://doi.org/10.1093/bioinformatics/btr330

Davey, J. W., & Blaxter, M. L. (2010). RADSeq: Next-generation population genetics. Briefings in Functional Genomics, 9(5–6), 416–423. https://doi.org/10.1093/bfgp/elq031

Davidson, N. M., Hawkins, A. D. K., & Oshlack, A. (2017). SuperTranscripts: A data driven reference for analysis and visualisation of transcriptomes. Genome Biology, 18(1), 148. https://doi.org/10.1186/s13059-017-1284-1

Davidson, N. M., & Oshlack, A. (2014). Corset: Enabling differential gene expression analysis for de novoassembled transcriptomes. Genome Biology, 15(7), 410. https://doi.org/10.1186/s13059-014-0410-6

Department of Fisheries. (1891). Report of the Department of Fisheries for the Year 1890. Ottawa. Retrieved from https://waves-vagues.dfo-mpo.gc.ca/Library/40758606_1890_pt.1.pdf.

Dirzo, R., Young, H. S., Galetti, M., Ceballos, G., Isaac, N. J. B., & Collen, B. (2014). Defaunation in the Anthropocene. Science, 345(6195), 401–406. https://doi.org/10.1126/science.1251817

Dobin, A., Davis, C. A., Schlesinger, F., Drenkow, J., Zaleski, C., Jha, S., Batut, P., Chaisson, M., Gingeras, T. R. (2013). STAR: Ultrafast universal RNA-seq aligner. Bioinformatics, 29(1), 15–21. https://doi.org/10.1093/bioinformatics/bts635

Ellison, C. E., Hall, C., Kowbel, D., Welch, J., Brem, R. B., Glass, N. L., & Taylor, J. W. (2011). Population genomics and local adaptation in wild isolates of a model microbial eukaryote. Proceedings of the National Academy of Sciences, 108(7), 2831–2836. https://doi.org/10.1073/pnas.1014971108

Falush, D., Stephens, M., & Pritchard, J. K. (2007). Inference of population structure using multilocus genotype data: Dominant markers and null alleles: Molecular Ecology Notes, 7(4), 574–578. https://doi.org/10.1111/j.1471-8286.2007.01758.x

Falush, D., Stephens, M., & Pritchard, J. K. (2003). Inference of Population Structure Using Multilocus Genotype Data: Linked Loci and Correlated Allele Frequencies. Genetics, 164(4), 1567 LP – 1587. Retrieved from http://www.genetics.org/content/164/4/1567.abstract

Fielder, D. G., Schaeffer, J. S., & Thomas, M. V. (2007). Environmental and ecological conditions surrounding the production of large year classes of walleye (*Sander vitreus*) in Saginaw Bay, Lake Huron. Journal of Great Lakes Research, 33(SUPPL. 1), 118–132. https://doi.org/10.3394/0380-1330(2007)33[118:EAECST]2.0.CO;2

Foll, M., & Gaggiotti, O. (2008). A genome-scan method to identify selected loci appropriate for both dominant and codominant markers: A Bayesian perspective. Genetics, 180(2), 977– 993. https://doi.org/10.1534/genetics.108.092221

Forney, J. L. (1976). Year-class formation in the walleye (*Stizostedion vitreum vitreum*) population of Oneida Lake, New York, 1966–73. Journal of the Fisheries Research Board of Canada, 33(4), 783–792. https://doi.org/10.1139/f76-096

Funk, W. C., McKay, J. K., Hohenlohe, P. A., & Allendorf, F. W. (2012). Harnessing genomics for delineating conservation units. Trends in Ecology & Evolution, 27(9), 489–496. https://doi.org/10.1016/j.tree.2012.05.012

Garrigan, D., & Hammer, M. F. (2006). Reconstructing human origins in the genomic era. Nature Reviews Genetics, 7(9), 669–680. https://doi.org/10.1038/nrg1941

Garrison, E., & Marth, G. (2012). Haplotype-based variant detection from short-read sequencing. ArXiv:1207.3907 [q-Bio]. Retrieved from http://arxiv.org/abs/1207.3907

Grabherr, M. G., Haas, B. J., Yassour, M., Levin, J. Z., Thompson, D. A., Amit, I., … Regev, A. (2011). Full-length transcriptome assembly from RNA-Seq data without a reference genome. Nature Biotechnology, 29(7), 644–652. https://doi.org/10.1038/nbt.1883

Hamidan, N., & Britton, J. R. (2015). Age and growth rates of the critically endangered fish *Garra ghorensis* can inform their conservation management. Aquatic Conservation: Marine and Freshwater Ecosystems, 25(1), 61–70.

Healy, T. M., & Schulte, P. M. (2019). Patterns of alternative splicing in response to cold acclimation in fish. The Journal of Experimental Biology, 222(5), jeb193516. https://doi.org/10.1242/jeb.193516

Henderson, B. A., Wong, J. L., & Nepszy, S. J. (1996). Reproduction of walleye in Lake Erie: Allocation of energy. Canadian Journal of Fisheries and Aquatic Sciences, 53(1), 127–133. https://doi.org/10.1139/f95-162

Hoffmann, A. A., & Willi, Y. (2008). Detecting genetic responses to environmental change. Nature Reviews Genetics, 9(6), 421–432. https://doi.org/10.1038/nrg2339

Hubisz, M. J., Falush, D., Stephens, M., & Pritchard, J. K. (2009). Inferring weak population structure with the assistance of sample group information. Molecular Ecology Resources, 9(5), 1322–1332. https://doi.org/10.1111/j.1755-0998.2009.02591.x

Hussey, P. J., Ketelaar, T., & Deeks, M. J. (2006). Control of the actin cytoskeleton in plant cell growth. Annual Review of Plant Biology, 57(1), 109–125. https://doi.org/10.1146/annurev.arplant.57.032905.105206

Jeffries, K. M., Connon, R. E., Verhille, C. E., Dabruzzi, T. F., Britton, M. T., Durbin-Johnson, B. P., & Fangue, N. A. (2019). Divergent transcriptomic signatures in response to salinity exposure in two populations of an estuarine fish. Evolutionary Applications, eva.12799. https://doi.org/10.1111/eva.12799

Johnston, I. A., Bower, N. I., & Macqueen, D. J. (2011). Growth and the regulation of myotomal muscle mass in teleost fish. Journal of Experimental Biology, 214(10), 1617–1628.

Johnston, T. A., Lysack, W., & Leggett, W. C. (2012). Abundance, growth, and life history characteristics of sympatric walleye (*Sander vitreus*) and sauger (*Sander canadensis*) in Lake Winnipeg, Manitoba. Journal of Great Lakes Research, 38, 35–46. https://doi.org/10.1016/j.jglr.2010.06.009

Jombart, T., Devillard, S., & Balloux, F. (2010). Discriminant analysis of principal components: A new method for the analysis of genetically structured populations. BMC Genetics, 11(1), 94. https://doi.org/10.1186/1471-2156-11-94

Jones, O. R., & Wang, J. (2010). COLONY: a program for parentage and sibship inference from multilocus genotype data. Molecular Ecology Resources, 10(3), 551–555. https://doi.org/10.1111/j.1755-0998.2009.02787.x

Kim, S., & Coulombe, P. A. (2010). Emerging role for the cytoskeleton as an organizer and regulator of translation. Nature Reviews Molecular Cell Biology, 11(1), 75–81. https://doi.org/10.1038/nrm2818

Kim, S., Wong, P., & Coulombe, P. A. (2006). A keratin cytoskeletal protein regulates protein synthesis and epithelial cell growth. Nature, 441(7091), 362–365. https://doi.org/10.1038/nature04659

Knaus, B. J., & Grünwald, N. J. (2017). VCFR: a package to manipulate and visualize variant call format data in R. Molecular Ecology Resources, 17(1), 44–53. https://doi.org/10.1111/1755-0998.12549

Knutsen, H., Olsen, E. M., Jorde, P. E., Espeland, S. H., André, C., & Stenseth, N. C. (2011). Are low but statistically significant levels of genetic differentiation in marine fishes ‘biologically meaningful’? A case study of coastal Atlantic cod: Biologically relevant genetic signals. Molecular Ecology, 20(4), 768–783. https://doi.org/10.1111/j.1365-294X.2010.04979.x

Kolosov, D., Bui, P., Chasiotis, H., & Kelly, S. P. (2013). Claudins in teleost fishes. Tissue Barriers, 1(3), e25391. https://doi.org/10.4161/tisb.25391

Kritzer, J. P., & Sale, P. F. (2004). Metapopulation ecology in the sea: From Levins’ model to marine ecology and fisheries science. Fish and Fisheries, 5(2), 131–140. https://doi.org/10.1111/j.1467-2979.2004.00131.x

Kuleshov, M. V., Jones, M. R., Rouillard, A. D., Fernandez, N. F., Duan, Q., Wang, Z., … Ma’ayan, A. (2016). Enrichr: A comprehensive gene set enrichment analysis web server 2016 update. Nucleic Acids Research, 44(W1), W90–W97. https://doi.org/10.1093/nar/gkw377

Lake Winnipeg Basin Indicator Series: Fish Populations. (2018). Manitoba Government, 6.

Lawson, D. J., van Dorp, L., & Falush, D. (2018). A tutorial on how not to over-interpret STRUCTURE and ADMIXTURE bar plots. Nature Communications, 9(1), 3258. https://doi.org/10.1038/s41467-018-05257-7

Li, H., Handsaker, B., Wysoker, A., Fennell, T., Ruan, J., Homer, N., … 1000 Genome Project Data Processing Subgroup. (2009). The sequence alignment/map format and SAMtools. Bioinformatics, 25(16), 2078–2079. https://doi.org/10.1093/bioinformatics/btp352

Li, R. (1995). Regulation of cortical actin cytoskeleton assembly during polarized cell growth in budding yeast. The Journal of Cell Biology, 128(4), 599–615. https://doi.org/10.1083/jcb.128.4.599

Liu, L., Ang, K. P., Elliott, J. A. K., Kent, M. P., Lien, S., MacDonald, D., & Boulding, E. G. (2017). A genome scan for selection signatures comparing farmed Atlantic salmon with two wild populations: Testing colocalization among outlier markers, candidate genes, and quantitative trait loci for production traits. Evolutionary Applications, 10(3), 276–296. https://doi.org/10.1111/eva.12450

Luu, K., Bazin, E., & Blum, M. G. B. (2017). pcadapt: An R package to perform genome scans for selection based on principal component analysis. Molecular Ecology Resources, 17(1), 67–77. https://doi.org/10.1111/1755-0998.12592

Manitoba Fishery Regulations, 1987. (1987). SOR/87-509. Retrieved from https://laws-lois.justice.gc.ca/eng/regulations/sor-87-509/index.html

Manitoba Sustainable Development. (2018). Manitoba Sustainable Development Annual Report *2017-*2018(p. 206). Retrieved from http://www.gov.mb.ca/finance/publications/annual.html

Manitoba Sustainable Development. (2019). Lake Winnipeg Measures to Enhance Sustainability. Retrieved from https://www.gov.mb.ca/sd/pubs/fish_wildlife/fish/quota_buyback_proposed_reg.pdf

Marshall, W. S., Breves, J. P., Doohan, E. M., Tipsmark, C. K., Kelly, S. P., Robertson, G. N., & Schulte, P. M. (2018). Clauding-10 isoform expression and cation selectivity change with salinity in salt-secreting epithelia of *Fundulus heteroclitus*. The Journal of Experimental Biology, 221(1), jeb168906. https://doi.org/10.1242/jeb.168906

Moles, M. D., Johnston, T. A., Robinson, B. W., Leggett, W. C., & Casselman, J. M. (2008). Is gonadal investment in walleye (*Sander vitreus*) dependent on body lipid reserves? A multipopulation comparative analysis. Canadian Journal of Fisheries and Aquatic Sciences, 65(4), 600–614. https://doi.org/10.1139/F07-186

Moles, M. D., Robinson, B. W., Johnston, T. A., Cunjak, R. A., Jardine, T. D., Casselman, J. M., & Leggett, W. C. (2010). Morphological and trophic differentiation of growth morphotypes of walleye (*Sander vitreus*) from Lake Winnipeg, Canada. Canadian Journal of Zoology, 88(10), 950–960. https://doi.org/10.1139/Z10-062

Murphy, M. D., & Crabtree, R. E. (2001). Changes in the age structure of nearshore adult red drum off west-central Florida related to recruitment and fishing mortality. North American Journal of Fisheries Management, 21(3), 671–678. https://doi.org/10.1577/1548-8675(2001)021<0671:CITASO>2.0.CO;2

Patro, R., Duggal, G., Love, M. I., Irizarry, R. A., & Kingsford, C. (2017). Salmon provides fast and bias-aware quantification of transcript expression. Nature Methods, 14(4), 417–419. https://doi.org/10.1038/nmeth.4197

Pritchard, J. K., Stephens, M., & Donnelly, P. (2000). Inference of population structure using multilocus genotype data. Genetics, 155(2), 945–959.

Pruyne, D., & Bretscher, A. (2000). Polarization of cell growth in yeast. Journal of Cell Science, 113(4), 571 LP – 585. Retrieved from http://jcs.biologists.org/content/113/4/571.abstract

R Core Team (2019). R: A language and environment for statistical computing. R Foundation for Statistical Computing, Vienna, Austria. URL https://www.R-project.org/.

Reid, A. J., Carlson, A. K., Creed, I. F., Eliason, E. J., Gell, P. A., Johnson, P. T. J., … Cooke, S. J. (2019). Emerging threats and persistent conservation challenges for freshwater biodiversity. Biological Reviews, 94(3), 849–873. https://doi.org/10.1111/brv.12480

Rideout, R. M., & Tomkiewicz, J. (2011). Skipped spawning in fishes: More common than you might think. Marine and Coastal Fisheries, 3(1), 176–189. https://doi.org/10.1080/19425120.2011.556943

Russello, M. A., Kirk, S. L., Frazer, K. K., & Askey, P. J. (2011). Detection of outlier loci and their utility for fisheries management: Outlier loci for fisheries management. Evolutionary Applications, 5(1), 39–52. https://doi.org/10.1111/j.1752-4571.2011.00206.x

Sheppard, Katie T., Davoren, G. K., & Hann, B. J. (2015). Diet of walleye and sauger and morphological characteristics of their prey in Lake Winnipeg. Journal of Great Lakes Research, 41(3), 907–915. https://doi.org/10.1016/j.jglr.2015.05.006

Sheppard, K.T., Hann, B. J., & Davoren, G. K. (2018). Growth rate and condition of walleye (*Sander vitreus*), sauger (*Sander canadensis*), and dwarf walleye in a large Canadian lake. Canadian Journal of Zoology, 96(7), 739–747. https://doi.org/10.1139/cjz-2017-0276

State of Lake Winnipeg: 1999 to 2007. (2011). Environment Canada. Retrieved from http://publications.gc.ca/pub?id=9.652328&sl=0

Stepien, C. A., Murphy, D. J., Lohner, R. N., Sepulveda-Villet, O. J., & Haponski, A. E. (2009). Signatures of vicariance, postglacial dispersal and spawning philopatry: population genetics of the walleye *Sander vitreus*. Molecular Ecology, 18(16), 3411–3428.

Stepien, C. A., Snyder, M. R., & Knight, C. T. (2018). Genetic divergence of nearby walleye spawning groups in central Lake Erie: implications for management. North American Journal of Fisheries Management, 38(4), 783–793.

Sveen, L. R., Timmerhaus, G., Torgersen, J. S., Ytteborg, E., Jørgensen, S. M., Handeland, S., … Takle, H. (2016). Impact of fish density and specific water flow on skin properties in Atlantic salmon (*Salmo salar L*.) post-smolts. Aquaculture, 464, 629–637. https://doi.org/10.1016/j.aquaculture.2016.08.012

Vähä, J.-P., Erkinaro, J., Niemelä, E., & Primmer, C. R. (2007). Life-history and habitat features influence the within-river genetic structure of Atlantic salmon. Molecular Ecology, 16(13), 2638–2654. https://doi.org/10.1111/j.1365-294X.2007.03329.x

Van Der Maaten, L. J. P., & Hinton, G. E. (2008). Visualizing high-dimensional data using t-sne. Journal of Machine Learning Research, 9, 2579–2605. https://doi.org/10.1007/s10479-011-0841-3

Verta, J.-P., & Jones, F. C. (2019). Predominance of cis-regulatory changes in parallel expression divergence of sticklebacks. ELife, 8, e43785. https://doi.org/10.7554/eLife.43785.001

Wang, J. (2004). Sibship Reconstruction From Genetic Data With Typing Errors. Genetics, 166(4), 1963–1979. https://doi.org/10.1534/genetics.166.4.1963

Waples, R. S. (1998). Separating the wheat from the chaff: Patterns of genetic differentiation in high gene flow species. Journal of Heredity, 89(5), 438–450. https://doi.org/10.1093/jhered/89.5.438

Waples, Robin S., & Gaggiotti, O. (2006). What is a population? An empirical evaluation of some genetic methods for identifying the number of gene pools and their degree of connectivity: What is a population? Molecular Ecology, 15(6), 1419–1439. https://doi.org/10.1111/j.1365-294X.2006.02890.x

Wasteneys, G. O., & Galway, M. E. (2003). Remodeling the cytoskeleton for growth and form: An overview with some new views. Annual Review of Plant Biology, 54(1), 691–722. https://doi.org/10.1146/annurev.arplant.54.031902.134818

Watkinson, D. A., & Gillis, D. M. (2005). Stock discrimination of Lake Winnipeg walleye based on Fourier and wavelet description of scale outline signals. Fisheries Research, 72(2–3), 193–203. https://doi.org/10.1016/j.fishres.2004.11.002

Weir, B. S., & Cockerham, C. C. (1984). Estimating *F*-statistics for the analysis of population structure. Evolution, 38(6), 1358–1370. https://doi.org/10.1111/j.1558-5646.1984.tb05657.x

Whitlock, M. C., & Lotterhos, K. E. (2015). Reliable detection of loci responsible for local adaptation: Inference of a null model through trimming the distribution of *F* _ST_. The American Naturalist, 186(S1), S24–S36. https://doi.org/10.1086/682949

WWF (2016). Living Planet Report 2016: Risk and resilience in a new era. WWF International, Gland, Switzerland. Retrieved from http://awsassets.panda.org/downloads/lpr_living_planet_report_2016.pdf

WWF (2018). Living Planet Report 2018: Aiming higher (Eds M. Grooten and R. E. A. Almond). WWF International, Gland, Switzerland. Retrieved from http://assets.wwf.ca/downloads/lpr2018_full_report_spreads.pdf

Yan, J., Song, Z., Xu, Q., Kang, L., Zhu, C., Xing, S., … Sang, T. (2017). Population transcriptomic characterization of the genetic and expression variation of a candidate progenitor of *Miscanthus* energy crops. Molecular Ecology, 26(21), 5911–5922. https://doi.org/10.1111/mec.14338

Yang, R.-C. (1998). Estimating hierarchical *F*-statistics. Evolution, 52(4), 950–956. https://doi.org/10.1111/j.1558-5646.1998.tb01824.x

Zheng, X., Levine, D., Shen, J., Gogarten, S. M., Laurie, C., & Weir, B. S. (2012). A high-performance computing toolset for relatedness and principal component analysis of SNP data. Bioinformatics, 28(24), 3326–3328. https://doi.org/10.1093/bioinformatics/bts606

