## Supplementary material for "Genomic signals found using RNA sequencing support conservation of walleye (*Sander vitreus*) in a large freshwater ecosystem": All Supplementary Materials

### Methods

*RNA extraction and sequencing*

Specific modifications to the RNeasy Plus Mini Prep Kit (QIAGEN, Venlo, Netherlands) extraction protocol are provided here. Samples were homogenized at 50 rotations per second for 6 min, using one metal bead in a TissueLyser LT (QIAGEN). Lysate was centrifuged for 5 min at 12,775 x g to force tissue to the bottoms of tubes, which we avoided pipetting in further steps. 350 µL of the homogenized lysate was transferred to the gDNA Eliminator spin column and centrifuged for 60 s, the flow-through was added 350 µL of 70% ethanol. 700 µL of the sample were transferred to a RNeasy spin column. All subsequent centrifuge steps were performed for a minimum of 60 s at 12,775 x g. During the Buffer RW1 and Buffer RPE wash steps, tubes were inverted prior to centrifuging to wash the upper portions of the spin columns. After centrifuging, while discarding flow-through from the wash buffers, collection tubes were dabbed dry on paper towels prior to the next step. RNase-free H2O was heated to 60 °C and 30 µL was used for elutions, and incubated on the spin column membrane for at least 1 min prior to the final centrifugation at 12,775 x g.

*RNA SNP Calling*

The protocol used for following the SuperTranscripts pipeline (Davidson et al. 2017) is provided here. To call single nucleotide polymorphisms (SNPs) from aligned reads, Picard version 2.18.9 (Broad Institute 2018) was used to add read groups to aligned files, sort reads by coordinate, index read files, then hard-clipped portions of a read that overlapped with regions outside of a gene. We used FreeBayes version 1.2.0, a haplotype-based program, to call SNPs from the processed file (Garrison & Marth 2012). An index of the transcriptome was created using Samtools version 1.9 (Li et al. 2009), then on a computing cluster (https://www.westgrid.ca/) the maximum number of open file descriptors was set to 2024 (ulimit -n 2024) and the maximum stack size to 102400 (ulimit -s 102400) prior to running FreeBayes in parallel over 48 threads with default options. Scripts used in the analyses for this manuscript are provided at https://github.com/BioMatt/Walleye_RNAseq.

**Tables & Figures**

Supplementary Table 1. Information for individual Lake Winnipeg walleye (*Sander vitreus*) collected in this study. Sample ID represents sample identifiers used in bioinformatics and analyses. Location represents spawning sites at which fish were collected, with the Red River in the south basin, Matheson Island in the channel connecting two basins, and the Dauphin River representing the north basin. Sex as identified during sub-lethal sampling is provided, along with year collected. The number of reads are total reads for each sample sequenced on an Illumina NovaSeq 6000 (Illumina, San Diego, California, USA), and % uniquely mapped reads is the percent of reads sequenced mapped Star to the Lace-constructed transcriptome, and thus used in subsequent analyses. Adegenet was used to generate 2- and 3-cluster population reassignments.

| Sample ID | Location | Sex | Year Collected | Number of Reads | % Reads Uniquely Mapped | 2 Clusters | 3 Clusters |
|---|---|---|---|---|---|---|---|
| Wall-002 | Red River | F | 2017 | 44,434,267 | 81.71 | 1 | 3 |
| Wall-005 | Red River | F | 2017 | 43,257,271 | 81.29 | 1 | 3 |
| Wall-006 | Red River | F | 2017 | 48,288,672 | 81.03 | 1 | 3 |
| Wall-008 | Red River | F | 2017 | 52,313,120 | 80.69 | 1 | 3 |
| Wall-011 | Red River | F | 2017 | 48,682,209 | 81.33 | 1 | 3 |
| Wall-016 | Red River | F | 2017 | 47,412,164 | 81.31 | 1 | 3 |
| Wall-019 | Red River | F | 2017 | 38,206,953 | 80.60 | 1 | 3 |
| Wall-020 | Red River | F | 2017 | 58,283,154 | 80.91 | 1 | 3 |
| Wall-171 | Matheson Island | F | 2017 | 54,846,547 | 82.40 | 2 | 3 |
| Wall-172 | Matheson Island | F | 2017 | 47,903,022 | 81.80 | 1 | 3 |
| Wall-173 | Matheson Island | F | 2017 | 48,351,276 | 82.07 | 1 | 3 |
| Wall-174 | Matheson Island | F | 2017 | 55,749,392 | 81.47 | 1 | 3 |
| Wall-175 | Matheson Island | F | 2017 | 47,721,630 | 81.78 | 1 | 3 |
| Wall-176 | Matheson Island | F | 2017 | 46,802,684 | 82.15 | 1 | 3 |
| Wall-177 | Matheson Island | F | 2017 | 41,243,681 | 82.00 | 1 | 3 |
| Wall-178 | Matheson Island | F | 2017 | 39,125,672 | 81.39 | 2 | 3 |
| Wall-182 | Dauphin River | F | 2017 | 40,569,210 | 81.88 | 2 | 2 |

| | | | | | | | |
|---|---|---|---|---|---|---|---|
| Wall-184 | Dauphin River | F | 2017 | 43,602,697 | 81.08 | 1 | 3 |
| Wall-185 | Dauphin River | F | 2017 | 39,405,325 | 82.04 | 2 | 2 |
| Wall-187 | Dauphin River | F | 2017 | 46,263,666 | 81.34 | 1 | 3 |
| Wall-188 | Dauphin River | F | 2017 | 44,367,660 | 82.12 | 2 | 2 |
| Wall-193 | Dauphin River | F | 2017 | 41,158,198 | 81.47 | 1 | 3 |
| Wall-196 | Dauphin River | F | 2017 | 46,924,282 | 81.53 | 2 | 2 |
| Wall-197 | Dauphin River | F | 2017 | 39,784,120 | 81.75 | 2 | 2 |
| Wall-210 | Red River | F | 2018 | 44,613,544 | 81.76 | 1 | 1 |
| Wall-212 | Red River | F | 2018 | 44,117,274 | 81.99 | 1 | 1 |
| Wall-213 | Red River | F | 2018 | 43,396,269 | 82.50 | 1 | 1 |
| Wall-214 | Red River | F | 2018 | 52,719,989 | 81.76 | 1 | 1 |
| Wall-215 | Red River | F | 2018 | 44,762,039 | 81.71 | 1 | 1 |
| Wall-216 | Red River | F | 2018 | 36,008,614 | 81.31 | 1 | 1 |
| Wall-217 | Red River | F | 2018 | 48,005,006 | 81.55 | 1 | 1 |
| Wall-218 | Red River | F | 2018 | 44,721,946 | 81.36 | 1 | 1 |
| Wall-290 | Matheson Island | F | 2018 | 48,612,467 | 81.44 | 1 | 1 |
| Wall-292 | Matheson Island | F | 2018 | 42,865,563 | 81.46 | 1 | 1 |
| Wall-294 | Matheson Island | F | 2018 | 49,734,421 | 81.36 | 1 | 1 |
| Wall-295 | Matheson Island | F | 2018 | 46,729,515 | 81.87 | 1 | 1 |
| Wall-296 | Matheson Island | UN | 2018 | 39,870,540 | 81.08 | 1 | 1 |
| Wall-297 | Matheson Island | F | 2018 | 36,812,535 | 80.63 | 1 | 1 |
| Wall-301 | Matheson Island | F | 2018 | 50,070,079 | 80.58 | 1 | 1 |
| Wall-302 | Matheson Island | F | 2018 | 44,648,079 | 81.38 | 1 | 1 |
| Wall-343 | Dauphin River | UN | 2018 | 42,217,434 | 81.45 | 1 | 1 |
| Wall-344 | Dauphin River | F | 2018 | 50,429,946 | 80.99 | 2 | 2 |
| Wall-345 | Dauphin River | F | 2018 | 38,506,818 | 82.02 | 2 | 2 |
| Wall-349 | Dauphin River | UN | 2018 | 41,990,932 | 80.54 | 2 | 2 |
| Wall-350 | Dauphin River | UN | 2018 | 40,341,008 | 81.33 | 2 | 2 |
| Wall-353 | Dauphin River | F | 2018 | 44,956,582 | 79.62 | 1 | 1 |
| Wall-354 | Dauphin River | F | 2018 | 37,591,445 | 81.43 | 2 | 2 |
| Wall-355 | Dauphin River | F | 2018 | 52,407,394 | 81.49 | 1 | 1 |

Supplementary Table 2. Weir & Cockerham's Pairwise $F_{ST}$ calculated with hierfstat between the Red River in the South Basin, Matheson Island in the channel, and Dauphin River in the North Basin for all 48 walleye (*Sander vitreus*) sampled in both 2017 and 2018, with a data set of 38,732 SNPs that do not show a strong year effect (between-year $F_{ST} \leq 0.01$ for all SNPs used here). 95% confidence intervals are provided in parentheses.

| | **Red River (South Basin)** | **Matheson Island (channel)** | **Dauphin River (North Basin)** |
|---|---|---|---|
| **Red River** | - | 0.0018 (0.0013 - 0.0022) | 0.0072 (0.0067 - 0.0078) |
| **Matheson Island** | | - | 0.0050 (0.0044 - 0.0055) |
| **Dauphin River** | | | - |

Supplementary Table 3. Gene ontology terms found using EnrichR, which represent genes that vary along a latitudinal gradient in Lake Winnipeg (PC1 in Figure 4). These terms represent 120 uniquely annotated genes with 386 outlier SNPs among them found with pcadapt ($q < 0.05$). Benjamini-Hochberg adjusted *p*-values and the number of genes represented in each term are provided. This analysis was performed using a set of 222,634 single nucleotide polymorphisms (SNPs) not filtered for Hardy-Weinberg Equilibrium or pruned for linkage disequilibrium, unlike the putatively neutral set of SNPs used for population structure analyses.

| **Gene Ontology Term** | **Adjusted *p*-value** | **Number of Genes** |
|---|---|---|
| purine ribonucleoside triphosphate binding (GO:0035639) | 0.017 | 10 |
| ATP binding (GO:0005524) | 0.017 | 8 |
| adenyl ribonucleotide binding (GO:0032559) | 0.021 | 8 |
| hydrogen-exporting ATPase activity, phosphorylative mechanism (GO:0008553) | 0.084 | 2 |

Supplementary Figure 1. Bayesian Information Criterion (BIC) values versus number of clusters found when using Adegenet to explore population reassignment in all 48 walleye (*Sander vitreus*) collected in 2017 and 2018, over three sites representing a latitudinal gradient in Lake Winnipeg. These values were found using 52,372 Hardy-Weinberg Equilibrium filtered and linkage disequilibrium pruned, putatively neutral single nucleotide polymorphisms.

Supplementary Figure 2. Principal Components Analysis (PCA) and t-SNE plots using 52,37 putatively neutral and all 222,634 high-quality SNPs performed using Adegenet and Rtsne, respectively. These analyses were performed on all 48 walleye (*Sander vitreus*) sampled from Lake Winnipeg, Manitoba, Canada, used in this study. Color shows site collected (red for Red River in the South Basin, yellow Matheson Island in the channel, and blue Dauphin River in the North Basin), circles showing walleye collected in 2017, and triangles showing walleye collected in 2018. A) Represents the PCA using neutral SNPs, B) represents a t-SNE using neutral SNPs, C) represents a PCA using all high-quality SNPs, and D) represents a t-SNE using all high-quality SNPs.

Supplementary Figure 3. PCA and t-SNE plots using 38,732 SNPs that do not show a strong year effect (between-year $F_{ST} \leq 0.01$ for all SNPs used here) performed using Adegenet and Rtsne, respectively. These analyses were performed on all 48 walleye (*Sander vitreus*) used in this study. Color shows site collected (red for Red River in the South Basin, yellow Matheson Island in the channel, and blue Dauphin River in the North Basin), circles showing walleye collected in 2017, and triangles showing walleye collected in 2018. A) Represents the PCA, and B) represents the t-SNE.

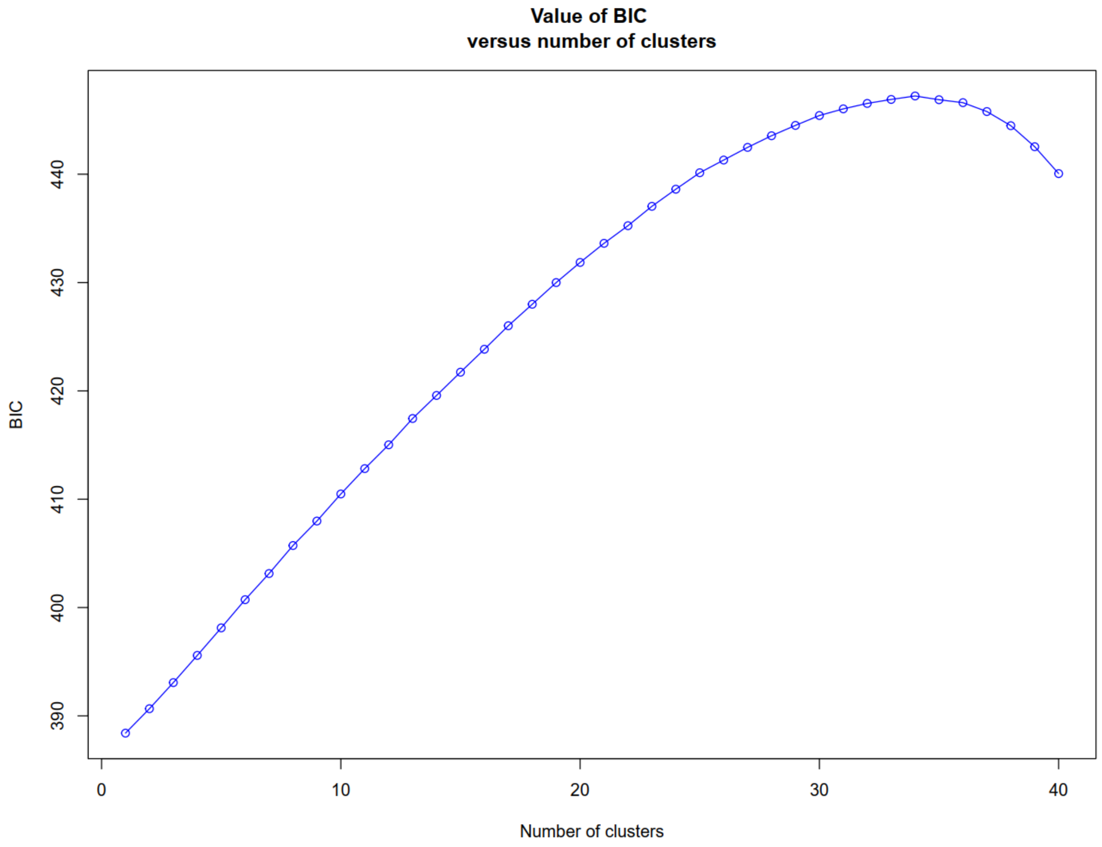
Value of BIC
versus number of clusters
BIC
390
400
410
420
430
440
0
10
20
30
40
Number of clusters

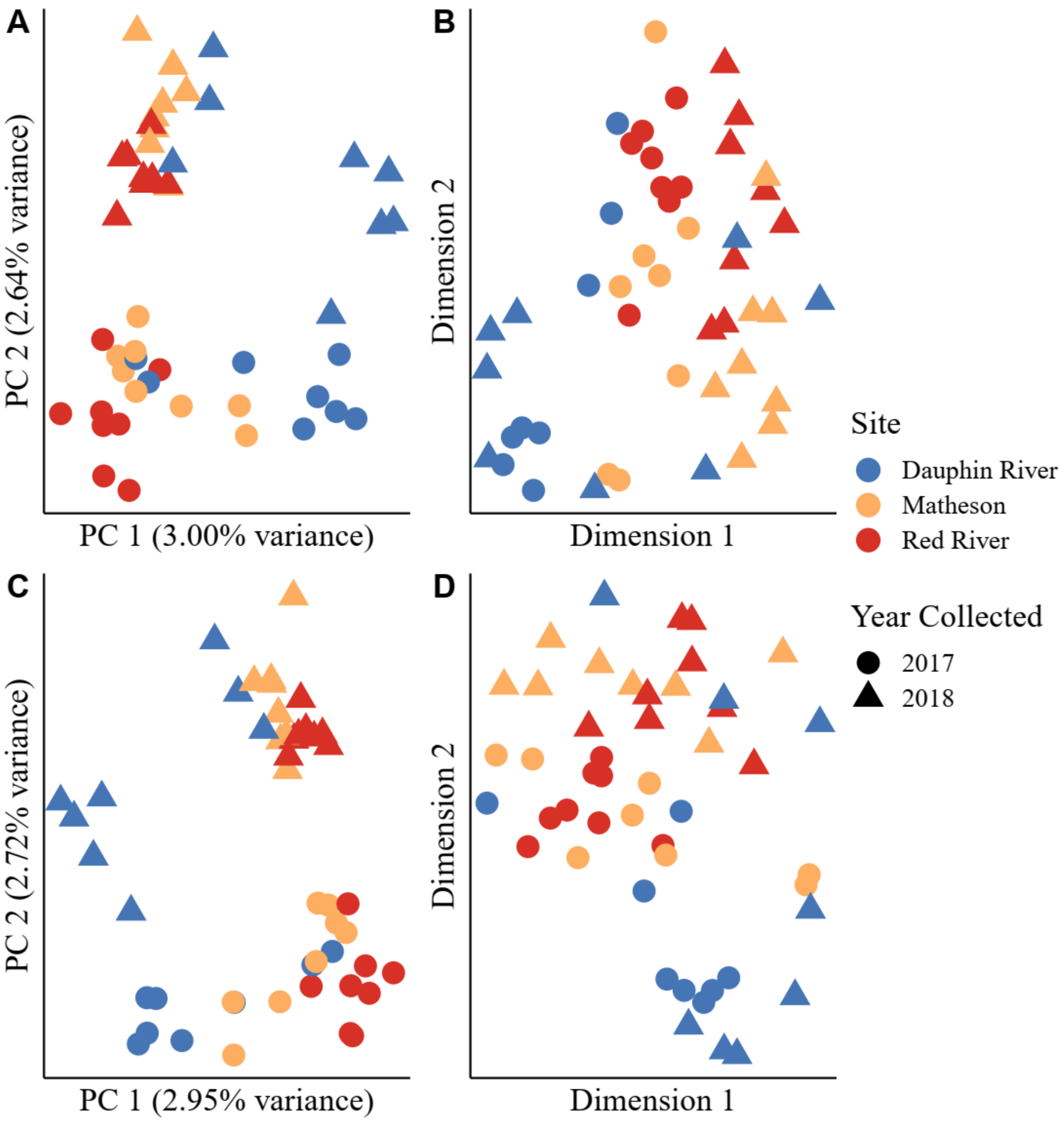
A
PC 2 (2.64% variance)
PC 1 (3.00% variance)
B
Dimension 2
Dimension 1
C
PC 2 (2.72% variance)
PC 1 (2.95% variance)
D
Dimension 2
Dimension 1
Site
Dauphin River
Matheson
Red River
Year Collected
2017
2018

124 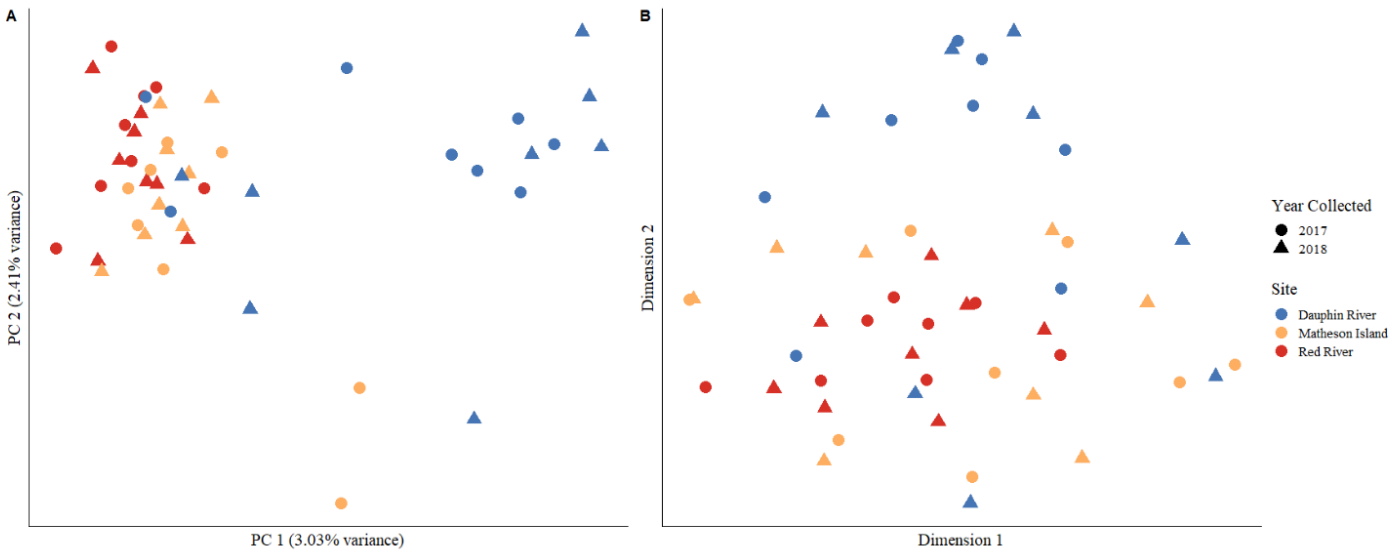
A
PC 2 (2.41% variance)
PC 1 (3.03% variance)
B
Dimension 2
Dimension 1
Year Collected
2017
2018
Site
Dauphin River
Matheson Island
Red River
